# FAN1 nuclease activity affects CAG expansion and age at onset of Huntington’s disease

**DOI:** 10.1101/2021.04.13.439716

**Authors:** Branduff McAllister, Jasmine Donaldson, Caroline S. Binda, Sophie Powell, Uroosa Chughtai, Gareth Edwards, Joseph Stone, Sergey Lobanov, Linda Elliston, Laura-Nadine Schuhmacher, Elliott Rees, Georgina Menzies, Marc Ciosi, Alastair Maxwell, Michael J. Chao, Eun Pyo Hong, Diane Lucente, Vanessa Wheeler, Jong-Min Lee, Marcy E. MacDonald, Jeffrey D. Long, Elizabeth H. Aylward, G. Bernhard Landwehrmeyer, Anne E. Rosser, REGISTRY Investigators of the European Huntington’s disease network, Jane S. Paulsen, PREDICT-HD Investigators of the Huntington Study Group, Nigel M. Williams, James F. Gusella, Darren G. Monckton, Nicholas D. Allen, Peter Holmans, Lesley Jones, Thomas H. Massey

## Abstract

The age at onset of motor symptoms in Huntington’s disease (HD) is driven by *HTT* CAG repeat length but modified by other genes. We used exome sequencing of 683 HD patients with extremes of onset or phenotype relative to CAG length to identify rare variants associated with clinical effect. We identified damaging coding variants in candidate modifier genes from prior genome-wide association studies associated with altered HD onset or severity. Variants in FAN1 clustered in its DNA-binding and nuclease domains and were associated predominantly with earlier onset HD. Nuclease activities of these variants correlated with residual age at motor onset of HD. Mutating endogenous FAN1 to a nuclease-inactive form in an induced pluripotent stem cell model of HD led to rates of CAG expansion comparable to those observed with complete *FAN1* knock out. Together, these data implicate FAN1 nuclease activity in slowing somatic repeat expansion and hence onset of HD.

## Introduction

Huntington’s disease (HD) is the most common monogenic degenerative disorder of the central nervous system. It is caused by an expanded CAG repeat of at least 36 trinucleotides in exon 1 of the *HTT* gene, with the expanded tract being both necessary and sufficient for disease (The Huntington’s Disease Collaborative Research Group, 1993; Lee et al., 2012a). Neuronal loss is initially most prominent in the striatum but becomes widespread in the brain, leading to a progressive movement disorder alongside functionally debilitating neuropsychiatric and cognitive decline. There is currently no disease-modifying treatment and premature death typically occurs 10 to 30 years after symptom onset (Bates et al., 2015).

Longer inherited CAG repeat tracts are associated with earlier onset of disease manifestations but they only account for ∼ 50% of the variation in onset observed (Andrew et al., 1993; Duyao et al., 1993; Snell et al., 1993; Lee et al., 2012a). Approximately 40% of the residual age at onset is heritable (Wexler et al., 2004), powering genome-wide association studies (GWAS) which have identified genetic variants that modify disease course (GeM-HD Consortium, 2015, 2019; Moss et al., 2017). The most recent Genetic Modifiers of HD (GeM-HD) GWAS identified 21 independent signals at 14 genomic loci associated with altered motor onset after accounting for inherited pure CAG length (GeM-HD Consortium, 2019). Candidate genes at these loci include *FAN1*, *MSH3*, *MLH1, PMS1, PMS2* and *LIG1*, all of which function in DNA repair. The most significant locus, on chromosome 15, includes *FAN1*, a gene encoding a 5’-exo/endo-nuclease involved in interstrand crosslink repair (Jin and Cho, 2017), and has four independent genome-wide signals (Kim et al., 2020). In addition, the pure CAG length of the pathogenic *HTT* repeat, rather than the length of the encoded polyglutamine, is most strongly associated with motor onset in individuals (Ciosi et al., 2019; GeM-HD Consortium, 2019; Wright et al., 2019). These data converge on the hypothesis that somatic (non-germline) expansion of the pathogenic inherited *HTT* CAG repeat in striatal neurons is a key driver of the rate of onset of HD in patients. In support of this model, somatic *HTT* CAG repeat expansions beyond the inherited repeat length in post-mortem human HD cortex correlate with age at motor onset (Swami et al., 2009), and some of the DNA repair variants implicated in modifying HD onset are associated with somatic CAG expansion in blood DNA from individuals with HD (Ciosi et al., 2019; Flower et al., 2019). The largest somatic CAG expansions are observed in the striatal neurons that degenerate earliest in HD in both human brain samples (Shelbourne et al., 2007; Gonitel et al., 2008) and mouse models of HD (Kovalenko et al., 2012). Furthermore, in mouse HD models, knockout of *Msh3*, *Mlh1* or *Mlh3* ablates somatic expansions of the CAG repeat (Dragileva et al., 2009; Pinto et al., 2013), whereas knockout of *Fan1* increases expansions (Loupe et al., 2020), in agreement with directions of effect predicted by human genetics.

GWAS can effectively identify common genetic variation associated with a disease or trait (Buniello et al., 2019). However, understanding pathogenic mechanisms through common variants can be difficult: over 90% of GWAS signals are non-coding and likely to either be tagging an actual functional coding variant or subtly modulating gene expression (Maurano et al., 2012). Even SNPs encoding protein sequence changes might not alter protein function if they are common in the population. The combined effect of all SNPs in a GWAS usually accounts for a maximum of one-third to two-thirds of trait heritability, with genome-wide significant SNPs contributing only a small percentage (Visscher et al., 2017). Some of the missing heritability is likely to be accounted for by rarer variants of larger effect size, such as loss-of-function or other damaging non-synonymous coding changes, that are not well captured by array-based imputation GWAS. Rare coding variants can give direct insight into molecular pathogenesis. For example, recent sequencing studies have identified disease-modifying roles for rare coding variants in schizophrenia (Xu et al., 2011; Fromer et al., 2014; Rees et al., 2020), Alzheimer’s disease (Bis et al., 2018; Raghavan et al., 2018), amyotrophic lateral sclerosis (van Rheenen et al, 2016), and diabetes mellitus (Flannick et al., 2019).

The putative HD modifier genes *FAN1*, *LIG1*, *MSH3*, *PMS1* and *PMS2* are all predicted to be tolerant of loss-of-function variation in the population (pLI < 0.02), supporting the possibility of uncovering rare modifier coding variants of large effect size in these genes (Karczewski et al., 2020). To this end, we sequenced the exomes of 683 HD patients with early onset/more severe or late onset/less severe symptoms relative to those predicted by pure CAG length alone, and looked for associations of coding variants with phenotype.

## Results

### Selection of HD study population with extremes of onset or phenotype

To maximise power to detect rare modifier variants in our sequencing cohort, we included individuals with extreme HD phenotypes from two independent longitudinal studies. First, we stratified 6086 participants from the retrospective Registry study by their residual age at motor onset, the difference between the actual age at motor onset and that predicted by pure CAG length alone (Langbehn et al., 2004), and selected ∼ 4% at each extreme for investigation (Figure 1A; 250 early onset, 250 late onset; ‘Registry-HD group’). Second, we selected participants from the prospective Predict-HD study (Paulsen et al., 2008) for investigation based on either extreme cognitive or motor phenotypes, or extreme predicted early or late onset, or both (Figure 1B-1D; ∼ 11% at each extreme; N=232/1069; ‘Predict-HD group’). CAG lengths used in initial sample selections came from standard PCR-fragment length assays that assume a canonical CAG repeat sequence.

**Figure 1.**
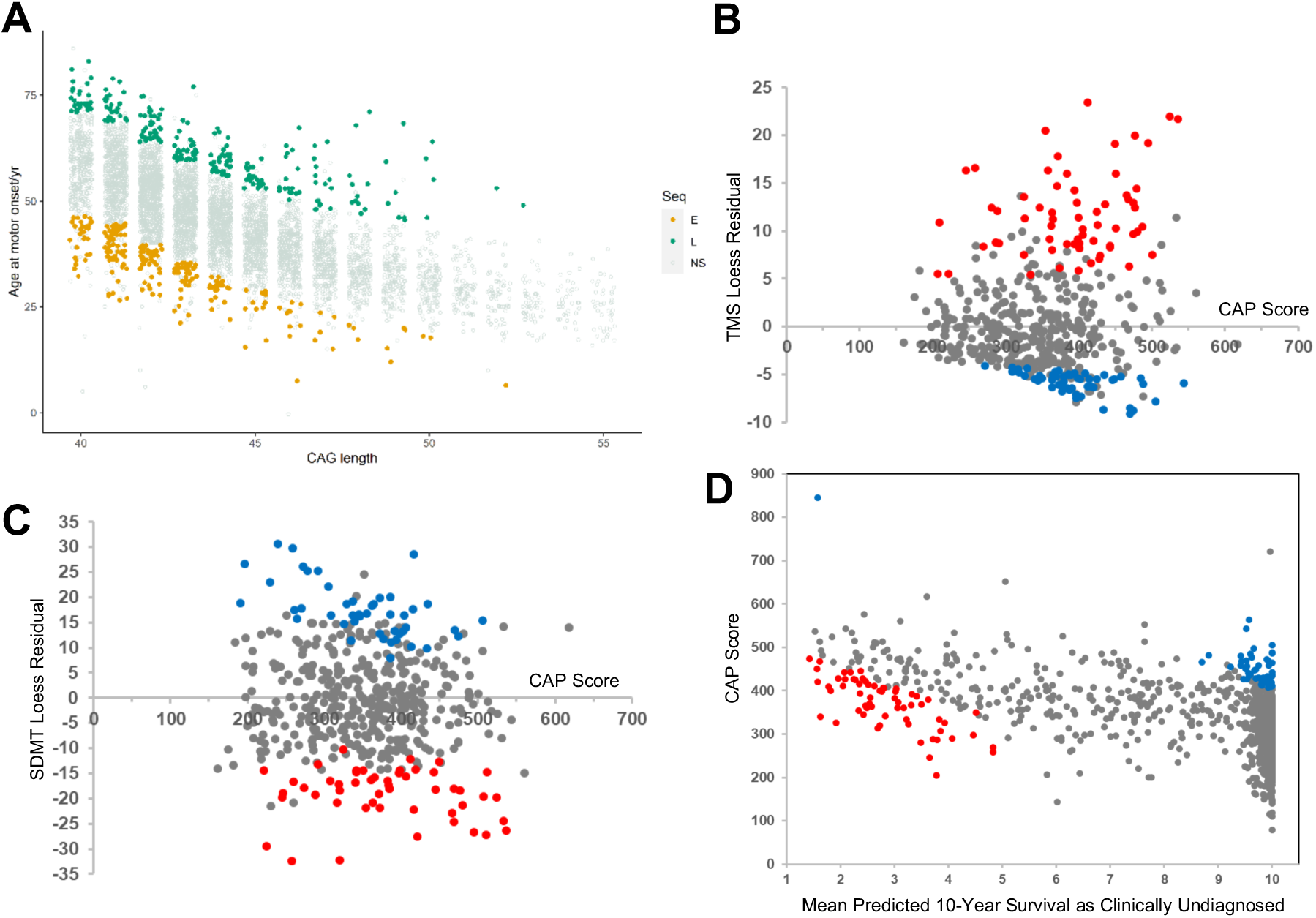
Selection of HD study population with extremes of onset or phenotype. (A) Registry-HD group. Age at motor onset against inherited CAG length for 6086 HD patients with 40-55 CAGs in the Registry study, using repeat lengths previously determined by PCR fragment length analysis. Individuals with very early (orange, N=250) or very late (green, N=250) motor onset given their inherited CAG length were selected for analysis (B - C) Predict-HD group, extremes of phenotype. Individuals with more severe (red dots) or less severe (blue dots) clinical phenotypes in the Predict-HD cohort were selected for analysis. Residuals from locally weighted smoothing (LOESS) were used to identify individuals using (B) total motor score (TMS; N=117) or (C) cognition score (symbol digit modalities test (SDMT); N=85), and are plotted against CAG age product (CAP) score to visualise age and CAG effects. Higher scores represent greater disease burden. (D) Predict-HD group, predicted early or late onset. A time-to-onset model was used to stratify the Predict-HD population and select a further cohort of predicted extreme early (red dots) or late (blue dots) onset individuals (N=119 selected).

### Non-canonical *HTT* CAG repeat sequences in expanded alleles are associated with altered clinical onset after correcting for CAG length

The polyglutamine-encoding pure CAG repeat in *HTT* exon 1 is typically followed by a glutamine-encoding CAACAG cassette and a polyproline-encoding CCG/CCA/CCT repeat (Figure 2, allele groups a and b). This region of *HTT* is highly polymorphic and can impact on disease phenotype (Ciosi et al., 2019; GeM-HD Consortium, 2019; Wright et al., 2019). To investigate associations of the precise repeat sequence with motor onset, we sequenced the *HTT* CAG repeat locus in the Registry-HD group using ultra-high-depth MiSeq sequencing (N=419 passing quality control) (Ciosi et al., 2019). Of these participants, 398 were in GeM-HD GWAS (GeM-HD Consortium, 2019). We identified 16 independent *HTT* trinucleotide repeat structures downstream of the pure CAG repeat, eight of which occurred exclusively on pathogenic *HTT* alleles with an expanded CAG repeat (Figure 2B). A canonical CAACAG followed the pure CAG tract in 94% of all alleles (Figure 2, allele groups a and b). The proportion of non-canonical glutamine-encoding repeats was enriched in alleles containing an expanded CAG repeat (41/419 (9.8%)) relative to those with an unexpanded repeat (9/419 (2.1%); *X*^2^ = 21.8, *p*= 3.1E-6; Figure 2, allele groups c - h). Within the expanded allele group, the distribution of specific non-canonical structures was highly skewed: alleles lacking CAACAG were observed in 21 individuals (5%, allele groups c and d), all of whom had earlier onset HD than predicted from sequenced CAG repeat length (mean residual = −10.2 years; 21/213 (9.9%) and 0/206 (0%) in early and late onset groups, respectively, *X*^2^ = 19.4, *p* = 1E-5). In contrast, alleles containing an extra CAACAG cassette, or extra CAA or CAC trinucleotides (allele groups e - h), were observed in 20 individuals and exclusively associated with later disease onset than predicted by sequenced CAG repeat length (mean residual = +10.4 years; 0/213 (0%) and 20/206 (9.7%) in early and late onset respectively, *X*^2^ = 19.6, *p* = 9E-6).

**Figure 2.**
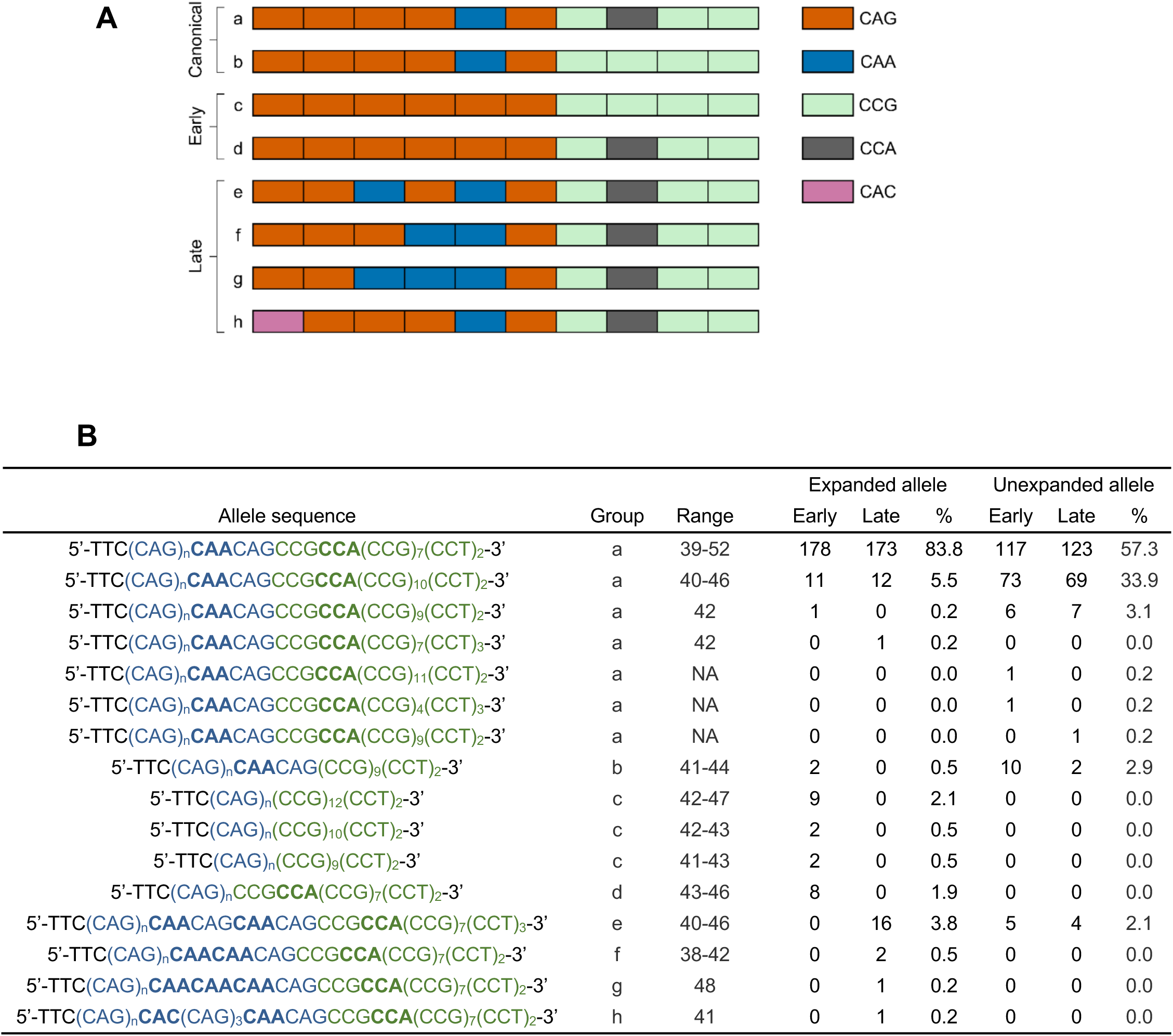
Non-canonical *HTT* CAG repeat sequences in expanded alleles are associated with altered clinical onset after correcting for pure CAG length. (A) Graphical overview of the 3’ end of the *HTT* exon 1 CAG repeat showing canonical (a - b) and non-canonical (c - h) trinucleotide arrangements identified by ultra-high-depth MiSeq sequencing in the Registry-HD cohort. (B) *HTT* CAG repeat allele sequences and counts from ultra-high-depth MiSeq sequencing of the Registry-HD cohort (N=419 passing quality control; N=213 with early onset relative to inherited pure CAG length; N=206 with late onset relative to inherited pure CAG length. Note that CAG lengths derived from MiSeq data). Allele counts for expanded (pathogenic) and unexpanded (wild-type) *HTT* alleles are shown, for both early and late onset groups. The glutamine-encoding repeat tract is in blue, the 3’ proline-encoding repeat tract in green. Allele groups refer to those illustrated in (A). Interrupting trinucleotides within the CAG and CCG tracts are highlighted in bold. Range refers to pure CAG lengths in expanded alleles, where applicable.

Much of the apparent effect of non-canonical glutamine-encoding sequences in *HTT* on age at HD onset has been considered spurious, attributable to mis-assignment of the true residual age-at-onset when CAG length calculations have assumed a canonical allele sequence (GeM-HD Consortium, 2019; Hong et al., 2021). To investigate in our sample, we used linear regression to quantify the variance in age at onset explained by polyglutamine tract length, pure CAG length, and the presence or absence of non-canonical sequences (Table S1; N=558; 463 Registry-HD, 95 Predict-HD with motor onset). Polyglutamine length accounted for 31.0% (p = 7.2E-47) of the variance in motor onset, which rose to 37.2% (p = 3.1E-58) when regressing on pure CAG length instead. However, non-canonical *HTT* CAG tract sequences in expanded alleles remained significantly associated with altered age at onset even after accounting for pure CAG tract length. Modelling the effects of CAG tracts lacking a CAACAG cassette increased R^2^ by 1.0% (p = 4.0E-3), equating to a ∼16% decrease in age at motor onset in individuals carrying this variant. Likewise, modelling the effects of extra CAACAG or CAA or CAC increased R^2^ by 1.4% (p = 2.0E-4), equating to a ∼26% increase in age at motor onset in individuals carrying one of these variants. Together these effects increased R^2^ by 2.1% (Table S1). The effects of non-canonical sequences on phenotype after accounting for CAG tract length were also observed in a larger, dichotomous sample using logistic regression with Firth’s correction (N=637; 421 Registry-HD, 216 Predict-HD; see below for details). Lack of CAACAG cassette was significantly associated with early/more severe phenotype (p=1.2E-7); extra CAACAG or CAA or CAC were significantly associated with late/less severe phenotype (p=1.5E-6). Overall, these data indicate that the precise sequence of the *HTT* glutamine-encoding repeat explains a significant, but small, proportion of the variance in HD onset not explained by CAG repeat length.

Finally, GeM-HD GWAS reported SNPs tagging haplotypes that contain non-canonical *HTT* allele sequences: rs764154313 tagged a pure CAG repeat lacking a CAACAG cassette (CAG)_n_(CCG)_12_(CCT)_2_ (Figure 2B, group c, top), and rs183415333 tagged an allele with an extra CAACAG cassette, (CAG)_n_(**CAACAG**)_2_CCGCCA(CCG)_7_(CCT)_3_ (Figure 2B, group e; (GeM-HD Consortium, 2019)). To estimate how effective SNP genotyping is for identifying non-canonical *HTT* alleles in our sample, we compared GWAS SNP data with our *HTT* sequencing results in those individuals with both measures (N=398). The minor allele of SNP rs764154313 tagged six of the nine (CAG)_n_(CCG)_12_(CCT)_2_ alleles but none of the other alleles lacking CAACAG (Figure 2B, group c - d). The minor allele of SNP rs183415333 tagged the (CAG)_n_(**CAACAG**)_2_CCGCCA(CCG)_7_(CCT)_3_ allele (Figure 2, group e) in 15/16 (93.8%) of individuals carrying this non-canonical repeat sequence in an expanded CAG repeat (but 0/9 carrying it in an unexpanded repeat). Therefore, SNP data identified 28.6% (6/21) of CAG repeats lacking CAACAG, and 75.0% (15/20) of CAG repeats with an extra CAACAG or CAA or CAC, suggesting that the prevalence of these non-canonical CAG alleles in HD is not fully captured by common variant analyses.

### Candidate gene analysis shows that rare coding variation in *FAN1* is associated with modified onset of HD

Using data from GeM-HD GWAS (GeM-HD Consortium, 2019), we calculated that all SNPs accounted for 25.3% (+/− 4.5%) of the residual age at motor onset. Given that approximately 40% of the residual age at onset is heritable, there remains significant missing heritability. Therefore we investigated whether rare protein coding variants could be modifiers of HD phenotype by sequencing the exomes of our cohort and calling sequence variants (N=683 after quality control; Figures S1-S3). Two groups of exomes were used for downstream analyses, adjusted for accurate CAG repeat lengths taken from sequencing data. First, a dichotomous group (N=637) divided into extremes of phenotype: early actual or predicted onset relative to inherited CAG repeat length, or more severe motor or cognitive phenotype (N=315; early/more severe group) and late actual or predicted onset relative to inherited CAG repeat length, or less severe motor or cognitive phenotype (N=322; late/less severe group). Second, a continuous phenotype group containing all those with known age at motor onset and hence calculable age-at-onset residual (N=558). We assessed the association between rare non-synonymous coding variants and clinical phenotype in the 13 candidate modifier genes (other than *HTT)* identified in GeM-HD GWAS (GeM-HD Consortium, 2019), using logistic regression in the dichotomous group and linear regression in the continuous group. Rare variants were defined as having a minor allele frequency (MAF) of less than 1%. Independent analyses were performed with three groups of variants: all non-synonymous variants; non-synonymous variants predicted to be damaging to protein function (CADD PHRED score ≥ 20 indicating in 1% predicted most damaging variants; NSD20); and loss-of-function variants such as nonsense, frameshift and splice donor/acceptor variants. *FAN1* showed a significant signal in the dichotomous non-synonymous analysis after multiple testing correction (p= 2.3E-4), and approached significance in the NSD20 analyses of both dichotomous and continuous groups (Table 1). Five other candidate genes had nominally significant associations in at least one analysis: *PMS1*, *MSH3*, *TCERG1*, *RRM2B* and *LIG1* (Table 1). These candidate genes were the most significant at their respective genomic loci when assessing all genes (Table S4). There were marked skews in the distribution of deleterious variants in *FAN1*, *PMS1* and *MSH3* between the early/more severe and late/less severe phenotype groups: those in *FAN1* were more often associated with early/more severe phenotype, whereas those in *PMS1* and *MSH3* were more often associated with late/less severe phenotype (Table S5). Rare predicted deleterious coding variants in *PMS1* occurred in 14 individuals with late/less severe phenotype including one loss-of-function frameshift variant (Table S6). Two individuals carried a predicted damaging variant in *PMS1* in the early/more severe phenotype group. Although the distribution of rare, deleterious *MSH3* variants was less skewed (eight in people with early onset, 14 in late onset), loss-of-function variants were exclusively found in seven people with late/less severe phenotype. These *MSH3* loss-of-function variants were either extremely rare or previously unreported, and included 4 splice acceptor variants and 3 truncations (Table S6).

**Table 1.** Candidate gene analysis shows that rare coding variation in FAN1 is associated with modified onset of HD. Optimal Sequence Kernel Association Tests (SKAT-O) for 13 candidate modifier genes from GeM-HD GWAS (GeM-HD Consortium, 2019). Gene-wide variant numbers were regressed on either dichotomous early/more severe or late/less severe phenotypes (N=637; logistic regression) or a continuous phenotype of residual age at motor onset using pure CAG counts from sequencing (N=558; linear regression). Only rare variants were included (MAF < 0.01). Three different variant groups were tested: all non-synonymous (NS), non-synonymous and predicted damaging to protein function (NSD; CADD PHRED score ≥ 20), and loss-of-function (LoF). Chromosomal loci from GeM-HD GWAS are indicated. Significant associations in bold (p < 6.4E-4, Bonferroni correction for 13 genes and 6 tests); nominally significant associations in italics (p < 0.05). See also Table S4.

| Chr | Gene | Dichotomous group (N=637) |  |  | Continuous group (N=558) |  |  |
| --- | --- | --- | --- | --- | --- | --- | --- |
|  |  | NS | NSD20 | LoF | NS | NSD20 | LoF |
| 2A | *PMS1* | 2.72E-02 | 2.65E-03 | 3.13E-01 | 2.62E-01 | 1.08E-01 | NA |
| 3A | *MLH1* | 1.72E-01 | 5.58E-02 | NA | 4.99E-01 | 7.45E-01 | NA |
| 5A | *DHFR* | 1.05E-01 | 9.83E-02 | NA | 1.54E-01 | 1.42E-01 | NA |
| 5A | *MSH3* | 5.96E-02 | 2.78E-01 | 9.51E-03 | 3.00E-01 | 6.35E-01 | 1.09E-01 |
| 5B | *TCERG1* | 1.25E-02 | 4.54E-01 | NA | 2.74E-03 | 2.76E-01 | NA |
| 7A | *PMS2* | 1.00E+00 | 9.00E-01 | 5.55E-01 | 5.59E-01 | 3.51E-01 | 1.33E-01 |
| 8A | *RRM2B* | 1.69E-01 | NA | NA | 2.26E-02 | NA | NA |
| 8A | *UBR5* | 7.47E-01 | 7.23E-01 | NA | 4.11E-01 | 3.88E-01 | NA |
| 11A | *CCDC82* | 5.27E-01 | NA | NA | 6.09E-01 | 9.01E-01 | NA |
| 11B | *SYT9* | 1.05E-01 | 1.13E-01 | NA | 4.20E-01 | 4.12E-01 | NA |
| 15A | *FAN1* | **2.32E-04** | 6.61E-04 | 2.17E-01 | 1.09E-03 | 9.06E-04 | 3.40E-01 |
| 16A | *GSG1L* | 9.05E-01 | 9.13E-01 | NA | 4.09E-01 | 4.35E-01 | NA |
| 19A | *LIG1* | 9.47E-02 | 7.07E-02 | 5.09E-01 | 3.63E-02 | 2.30E-02 | 5.16E-01 |

### Exome-wide association analysis of rare variants with phenotype highlights *FAN1*

We explored whether an exome-wide analysis of all genes with sufficient rare, damaging variation would highlight any new putative modifiers of HD phenotype. Analysis of the dichotomous and continuous phenotype groups using SKAT-O with correction for CAG repeat lengths did not show any significant exome-wide associations (Table S7). *FAN1* was the only gene with p< 1E-03 in both the dichotomous and continuous analyses. A candidate pathway analysis, using significant pathways (q<1E-03) from Gem-HD GWAS identified one significant pathway (GO:0042578; phosphoric ester hydrolase activity, containing *FAN1*; p= 9.7E-05) and several nominally significant pathways, suggesting damaging rare variation may be important in pathways previously associated with common variation (Table S8). For example: GO:0006281 DNA repair is significantly enriched for damaging rare variation in the dichotomous analysis (p=1.2E-3) and also for common variant association in GeM GWAS (p=5.4E-11).

Of note, SKAT-O analysis without correcting for pure CAG repeat lengths found one exome-wide significant signal in *NOP14* on chromosome 4, 130 kbp upstream of *HTT* (p= 8.3E-06, continuous analysis). We found that NOP14 R697C (4:2943419:G:A) is in strong linkage disequilibrium with the pathogenic (CAG)_n_(CAACAG)_2_ *HTT* allele (Figure 2, allele group e, R^2^ = 0.902), explaining this spurious association of *NOP14* variation with HD phenotype.

### Rare deleterious *FAN1* variants are associated with altered HD onset and are clustered in functional domains

*FAN1* was the most significant gene in both common variant modifier GWAS and our candidate exome analysis. In our cohort 6.8% (N=43/637) were heterozygous for at least one rare, non-synonymous, predicted damaging coding variant in *FAN1* (Figure 3A). Two individuals carried two rare, damaging *FAN1* variants, although these could not be phased. One individual carried a loss-of-function frameshift variant, ST186SX (15:31197095:G:A). Of those carrying damaging *FAN1* variants, significantly more had an early/more severe phenotype than a late/less severe phenotype (Odds ratio 3.43, 95% CI 1.66-7.09, p = 8.9E-04). We identified 28 different rare coding non-synonymous FAN1 variants, including six previously-unreported mutations (encoding K168N, P366R, D498N, D702E, L713I and R969L). Those associated mostly with early/more severe HD clustered in the DNA binding and nuclease domains of FAN1, while a small cluster of variants associated with late/less severe HD mapped to the protein-protein interaction domain (Figure 3B). The R377W and R507H variants detected in GeM-HD GWAS, and also found here, map on the DNA binding domain of FAN1 and are found mostly in individuals with early/more severe disease. Of the seven individuals with late/less severe HD carrying FAN1 R377W or R507H, five were also genotyped in GeM-HD GWAS (GeM-HD Consortium, 2019): one carried an extra CAACAG cassette in the expanded *HTT* repeat tract (allele e, Figure 2); one is homozygous and two heterozygous for the common neutral I219V variant in MLH1 (rs1799977) associated with slightly later onset HD; one had high predicted FAN1 expression and one low predicted *MSH3* expression, although these could not be phased. Finally, five extremely rare (MAF < 1E-6) damaging variants were identified in the C-terminal nuclease domain and were exclusively associated with early/more severe disease. These data suggest that FAN1 DNA binding and nuclease activities are important for its role in delaying onset of HD.

**Figure 3.**
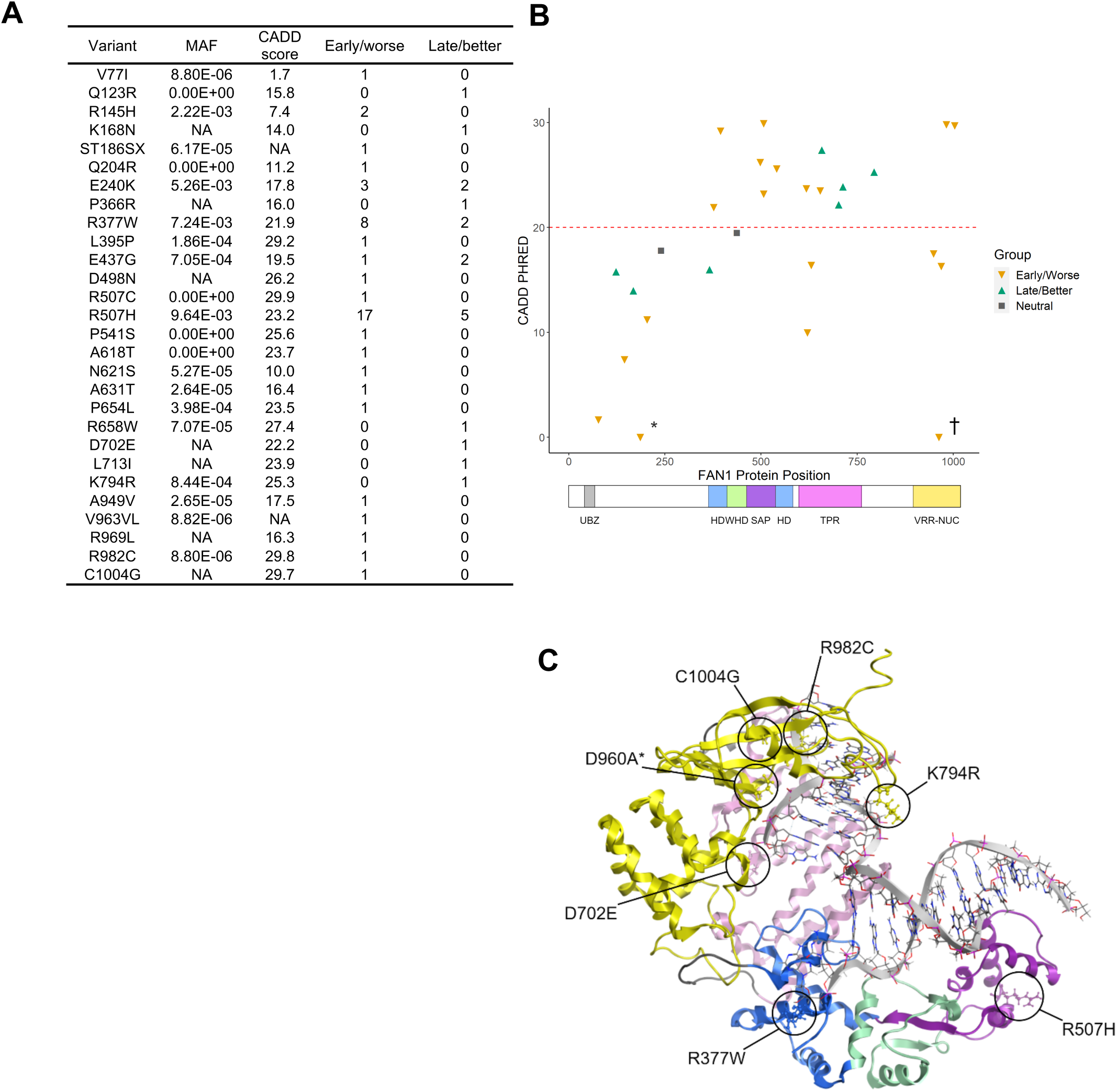
Rare deleterious FAN1 variants are associated with altered HD onset and are clustered in functional protein domains. (A) Rare, non-synonymous *FAN1* variants identified through exome sequencing in the dichotomous HD cohort (N=637), divided between early/more severe and late/less severe phenotype groups. MAF < 0.01. CADD score > 20 implies a variant is in the 1% predicted most damaging substitutions in the human genome. (B) FAN1 variants identified in individuals with HD, plotted by CADD score of predicted deleteriousness over a cartoon of FAN1 protein. Variants associated with early/more severe phenotype (orange triangles), late/less severe phenotype (green triangles), or neither phenotype group (grey squares, ‘neutral’) are shown. Two likely damaging singleton variants lack CADD scores and so are plotted as CADD=0. They are highlighted: loss-of-function (frameshift) variant T187SfsX3 (*) and in-frame insertion variant V936W964insL (†). FAN1 domain coordinates as published (Jin and Cho, 2017; UniProt Consortium, 2019). (C) Three-dimensional model highlighting FAN1 variants selected for downstream study. Note D960A (*) is a synthetic variant lacking nuclease activity, and not found in our patient population.

### The nuclease activity of FAN1 variants identified in individuals with HD correlates with residual age at onset of motor symptoms

The nuclease activity of FAN1 is required for its role in interstrand DNA crosslink (ICL) repair but has previously been shown to have no significant effect on CAG repeat expansion in a cellular model (Goold et al., 2019). We assessed whether four lymphoblastoid cell lines derived from individuals with HD heterozygous for the R507H FAN1 variant were as efficient in ICL repair as four age- and CAG length-matched controls with confirmed wild-type FAN1 alleles. Each R507H line was more sensitive to mitomycin C than its matched counterpart (Figure S4A) and, overall, the mean IC50 for the R507H lines was significantly lower than for the matched lines (Figure 4A; p=1.3E-03) suggesting that R507H is deleterious to FAN1 function, as previously suggested (Bastarache et al., 2018; Kim et al., 2020). Next, we selected six predicted deleterious FAN1 variants identified by exome sequencing for *in vitro* biochemical analysis of their nuclease activity. These variants (R377W, R507H, D702E, K794R, R982C and C1004G) as well as wild-type FAN1 and a known nuclease-inactive variant (D981A R982A) were expressed and partially purified from *E. coli* as NusA-His-tagged full-length proteins (Figure S4B-C (MacKay et al., 2010)). The flap endonuclease activities of wild-type and variant FAN1 proteins were assayed on canonical FAN1 substrates with short 5T 5’-flaps. Wild-type FAN1 converted 66% of 5’-flap substrate to product in a ten-minute reaction (Figure 4B, lane 1). The known nuclease-inactive double mutant lacked all nuclease activity, as expected (Figure 4B, lane 7). The predicted damaging variants all had reduced FAN1 activity compared to wild-type (Figure 4C), but variants associated mostly with early/more severe phenotype (R377W, R507H, R982C and C1004G) had much reduced activity compared with the two variants found in individuals with late/less severe phenotype (D702E and K794R). There was a significant correlation between mean residual age at onset for individuals harbouring each variant and nuclease activity of that variant (p= 2.5E-02), suggesting that FAN1 nuclease activity mediates its modifier role in HD.

**Figure 4.**
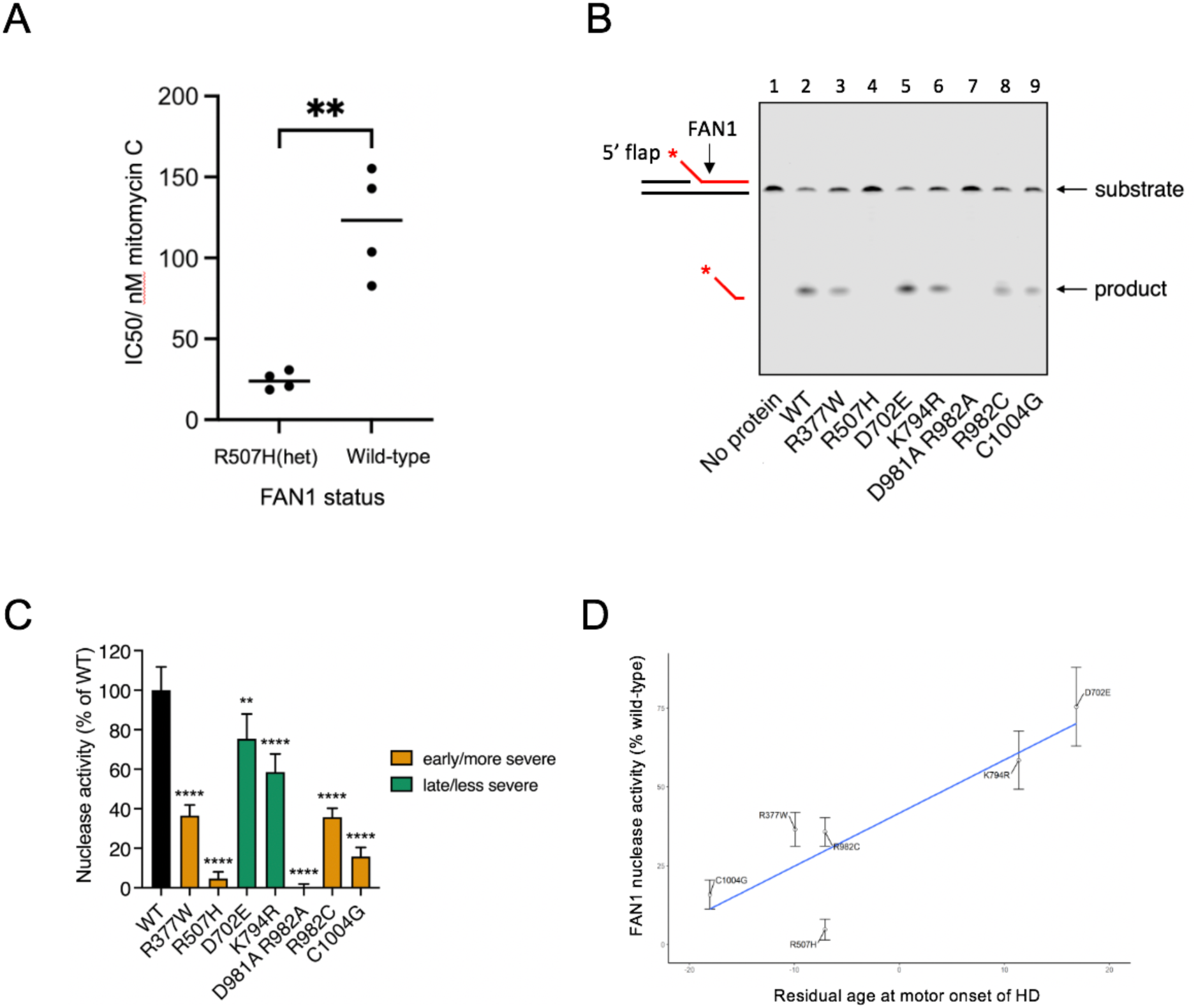
The nuclease activity of FAN1 variants identified in individuals with HD correlates with residual age at onset of motor symptoms. (A) Lymphoblastoid cell lines derived from patients carrying a heterozygous R507H FAN1 variant are significantly more sensitive to mitomycin C than age- and pure CAG length matched control HD lines with homozygous wild-type FAN1 (**p = 1.3E-03, two-tailed t-test, N=3 independent experiments) (B) Representative gel showing nuclease activity of FAN1 variants on 5’-flap DNA. FAN1 protein (10 nM) was incubated with 5’-flap DNA (5 nM) for 10 min at 37 °C in the presence of MnCl_2_. Reactions were denatured and analysed using 15% TBE-urea gel electrophoresis. (C) FAN1 variants identified in individuals with HD have significantly reduced nuclease activity compared with wild-type FAN1. Variants associated with early/more severe phenotypes (orange) had less nuclease activity than variants associated with late/less severe phenotypes (green). The nuclease-inactive D981A R982A FAN1 variant was used as a negative control. Activities of variants are normalised to wild-type FAN1 nuclease activity. (N=3, **p<1E-02, ****p<1E-04; One-way ANOVA; error bars, s.d.). (D) Graph of mean age at motor onset residual (using pure CAG length from sequencing) against FAN1 nuclease activity for 6 variants, normalised to wild-type FAN1 activity: R377W N=11; R507H N=17; D702E N=1; K794R N=1; R982C N=1; C1004 N=1. There was a significant correlation between average motor onset residual and *in vitro* nuclease activity (p = 2.5E-02). Three individuals had two FAN1 variants: C1004G & R507H; R982C & N621S; R377W & R507H. The rarer of the two variants was used in the analyses in each case. See also Figure S4.

### FAN1 slows the rate of *HTT* CAG repeat expansions in a nuclease-dependent manner in an induced pluripotent stem cell (iPSC) model of HD

To investigate the effect of *FAN1* knockout on *HTT* CAG repeat stability, we measured expansion rates in iPSCs derived from an individual with HD who inherited 109 CAGs (Q109 cells (HD iPSC Consortium, 2012)), with and without knocking out *FAN1* (Figures 5 & S5A-C). The *HTT* CAG repeat expanded significantly faster in cells lacking FAN1: each modal CAG unit increase occurred in 8.9 days in *FAN1*^-/-^ cells compared with 33.1 days in *FAN1*^WT/WT^ cells (p= 1.5E-5; Figure 5B-C). Expansion in differentiated neurons (Figure S5D) showed a similar effect although expansion rates were slower (Figure 5D). Each CAG unit increase occurred in 20.6 days in *FAN1*^-/-^ cells compared with 73.0 days in *FAN1*^WT/WT^ cells (p = 2.7E-5). Despite the slower rate of CAG expansion seen in neurons compared with iPSCs, the ratio of expansion rates *FAN1*^WT/WT^:*FAN1*^-/-^ is remarkably similar at 3.7x in iPSCs and 3.5x in neurons.

**Figure 5.**
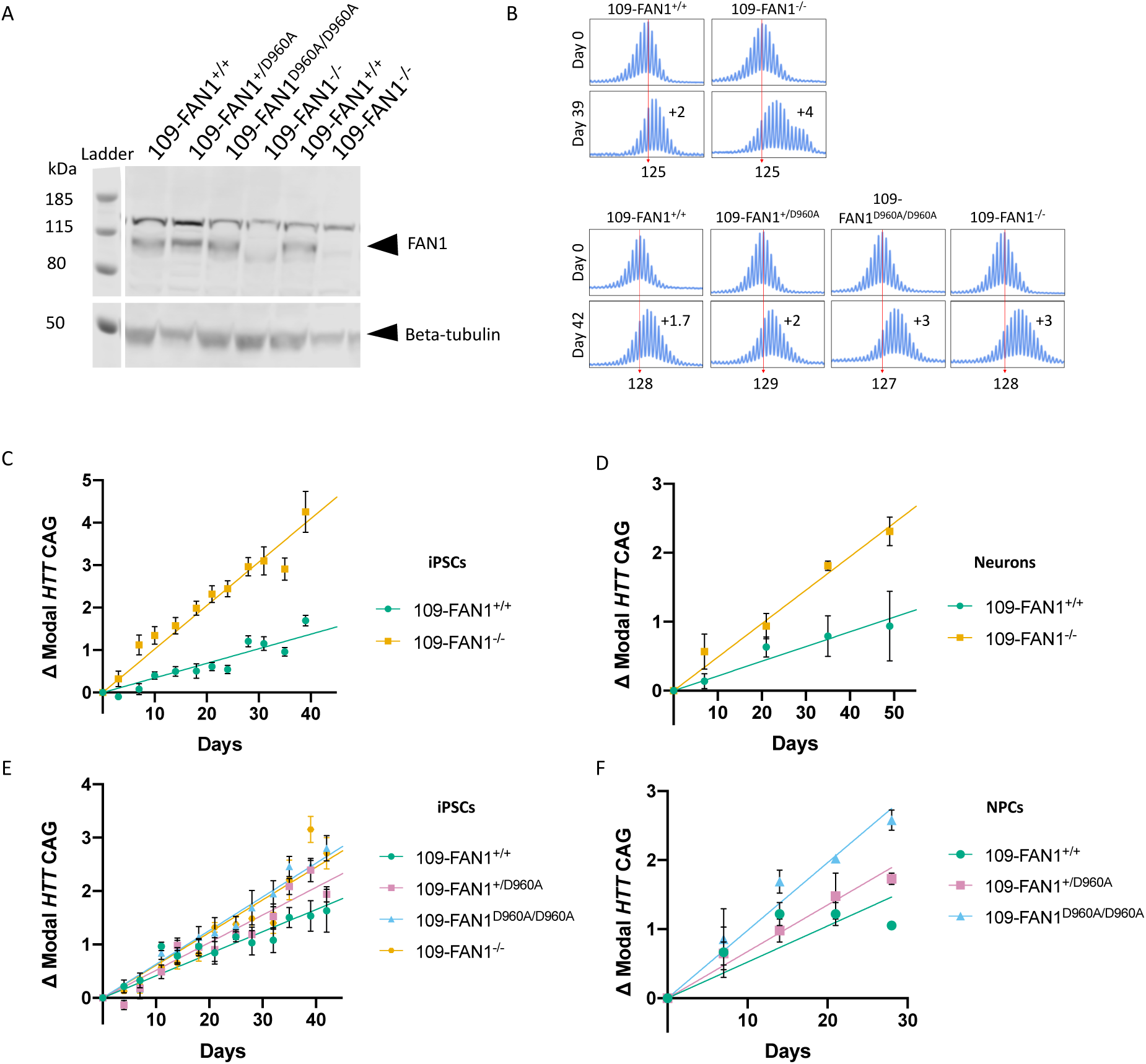
FAN1 slows the rate of *HTT* CAG repeat expansions in a nuclease-dependent manner in an induced pluripotent stem cell (iPSC) model of HD. (A) Western blot of FAN1 in Q109 HD iPSC lines. Parent Q109 line with wild-type FAN1 (lanes 1 and 5), Q109 lines with D960A variant (heterozygous (lane 2), homozygous (lane 3)), and Q109 FAN1 knockout lines (lanes 4 and 6). (B) Representative electropherograms of fluorescent PCR and capillary electrophoresis of the *HTT* CAG repeat in 109-*FAN1^+/+^*, 109-*FAN1^-/-^*, 109-*FAN1^+/D960A^* and 109-*FAN1^D960A/D960A^* iPSCs at baseline and after time in culture. The red dotted line indicates baseline *HTT* CAG repeat length. (C) 109-*FAN1^-/-^* iPSCs (N=6 clones) exhibit significantly faster *HTT* CAG repeat expansion rates than 109-*FAN1^+/+^* iPSCs (N=7 clones) (0.0661 CAG/day) (p= 1.5E-5). (D) Change in modal *HTT* CAG repeat in post-mitotic neurons generated from 109-*FAN1^+/+^* iPSCs (N=5) and 109-*FAN1^-/-^* iPSCs (N=4 clones). 109-*FAN1^-/-^* neurons demonstrate significantly faster rates of *HTT* CAG repeat expansion than 109-*FAN1^+/+^* neurons (p= 1.1E-2). (E) FAN1-D960A HD-iPSCs show dose-dependent increase in *HTT* CAG repeat expansion over time in culture. (N=3 clones per genotype, each cultured in triplicate wells). (F) D960A mutations enhance *HTT* CAG repeat expansions in a dose-dependent manner in neural precursor cells (NPCs) derived from 109-*FAN1^+/+^*, 109-*FAN1^+/D960A^* and 109-*FAN1^D960A/D960A^* iPSCs (N=1 clone per genotype, cultured in triplicate wells) Values expressed as mean ± SEM.

To investigate the importance of FAN1 nuclease activity in slowing *HTT* CAG repeat expansion, we used CRISPR/Cas9 to introduce a D960A FAN1 point mutation into Q109 cells (Figure S5E-G). D960A FAN1 lacks nuclease activity but retains wild-type DNA binding capacity (Kratz et al., 2010). Somatic instability of the *HTT* CAG tract was then assessed in *FAN1*^WT/WT^, *FAN1*^WT/D960A^, *FAN1*^D960A/D960A^, and *FAN1*^-/-^ Q109 cells (Figure 5E) and showed a nominally significant difference between genotypes (p=4.5E-02). Cells carrying homozygous or heterozygous D960A FAN1 had significantly greater rates of change in modal CAG length compared with wild-type cells (p= 2.3E-02), but there was no detectable difference between Q109 FAN1^WT/D960A^, FAN1^D960A/D960A^ and FAN1^-/-^ cells (p= 2.3E+01). Adding one CAG took 23.9 days in the FAN1^WT/WT^ cells, but 19.3 days in the FAN1^WT/D960A^, and 15.6 and 16.3 days in the FAN1^D960A/D960A^ and FAN1^-/-^ cells, respectively. These data suggest an additive effect of FAN1 nuclease activity on expansion.

We also saw a significant difference between genotypes in rates of expansion in a single clone of each FAN1 D960A genotype in neural precursor cells grown for 42 days (Figure 5F). A significant effect of genotype on rate of change in modal CAG was seen (p= 6.0E-03). Adding one CAG took 19.1 days in FAN1^WT/WT^ cells, 14.8 days in FAN1^WT/D960A^ cells and 10.8 days in FAN1^D960A/D960A^ cells. The rate of CAG expansion in FAN1^WT/D960A^ cells was significantly lower than that in FAN1^D960A/D960A^ cells (p= 6.0E-03), while the rate of CAG expansion in FAN1^WT/WT^ cells trends towards being lower than that in FAN1^WT/D960A^ cells (p= 6.6E-02). Fitting an additive model of D960A to CAG expansion rates increases significance over the general model (p= 3E-03), suggesting that there is a dose-dependent effect of the D960A mutation on CAG expansion rate, consistent with the effect seen in iPSCs.

## Discussion

Previous studies showed that inherited pathogenic *HTT* CAG repeat length has a stronger effect on age at onset of HD than polyglutamine length (Ciosi et al., 2019; GeM-HD Consortium, 2019; Wright et al., 2019). As pure CAG lengths are typically determined by a PCR-fragment length assay that assumes a canonical single CAACAG after the CAG tract, non-canonical glutamine-encoding *HTT* repeat sequences lead to mis-counting of pure CAG lengths. Use of these inaccurate counts leads to erroneous age-at-onset calculations in these individuals and explains most of the spurious association of non-canonical alleles with HD onset (GeM-HD Consortium, 2019). However, in our cohort of individuals with extremes of phenotype non-canonical glutamine-encoding *HTT* repeat sequences remained significantly associated with HD onset, even after adjusting for correct CAG repeat lengths determined by sequencing (Table S1). The difference in result between this study and GeM-HD GWAS might be because by sampling extremes of phenotype we have enriched for rare non-canonical sequences, and thus increased power to detect their phenotypic effects. Furthermore, we have shown here that *HTT* repeat locus sequencing identifies more non-canonical sequences than genotyping alone. These data indicate that it is not simply CAG length that influences disease onset; instead, the precise flanking sequence might also be important. Repeat tracts lacking a CAACAG cassette were exclusively associated with earlier onset disease, tracts with extra CAACAG or CAA or CAC exclusively with later onset disease. These sequence differences are most likely to mediate an effect through altered somatic expansion of the genomic pathogenic CAG repeat, with tracts lacking CAACAG expanding more easily and those with extra CAACAG or CAA or CAC expanding less readily. Alternative mechanisms, including effects on splicing or translation of RNA, and the influence of flanking repeats, cannot be ruled out.

Many of the candidate genes identified in modifier studies of HD encode proteins involved in DNA repair. Similar to the non-canonical glutamine-encoding repeat sequences in *HTT*, they are most likely to modify HD onset by modulating somatic expansion of the inherited expanded *HTT* CAG repeat in the brain (Kennedy et al., 2003), although an effect on global genomic stability cannot be entirely discounted. Somatic CAG expansion in post-mortem human brain is associated with earlier onset of HD (Swami et al., 2009), and is particularly marked in the striatal neurons that degenerate first in HD (Shelbourne et al., 2007; Gonitel et al., 2008), suggesting that it accelerates, or even causes, disease pathology in individuals carrying the HD mutation. Assessing candidate genes derived from GeM-HD GWAS (GeM-HD Consortium, 2019) we identified significant numbers of rare, predicted deleterious, coding variants in *FAN1* and nominally significant numbers in *PMS1*, *MSH3*, *TCERG1*, *RRM2B* and *LIG1* (Table 1). Furthermore, the directions of effect for rare damaging variants in *FAN1*, *PMS1* and *MSH3* were consistent with those observed in GeM-HD GWAS: *FAN1* variants were mostly associated with early/more severe phenotype, whereas *PMS1* variants and *MSH3* loss-of-function variants were mostly associated with late/less severe phenotype (Table S5). These data reinforce the conclusion that these genes are indeed driving the GWAS associations previously observed at their respective loci.

Finding rare modifier variants significantly associated with extremes of phenotype suggests that they might individually have large effect sizes. For example, in our sample, we found that lack of CAACAG after the CAG repeat was associated with ∼16% earlier onset than predicted by CAG length, while extra CAACAG or CAA or CAC was associated with ∼26% later onset than predicted (Table S1). However, effect sizes in the wider HD population cannot be determined from our cohort that was selected by extremes of phenotype. GeM-HD GWAS was able to assign effect sizes to moderately rare coding SNPs (MAF = 0.5 – 1.0%) in a large unselected HD population. For example, damaging coding variants in FAN1 had large effect sizes: each R507H allele was associated with 5.2 years earlier onset, and each R377W allele with 3.8 years earlier onset (GeM-HD Consortium, 2019). The extreme rarity of many of the deleterious coding variants we report here makes it difficult to estimate their true effect sizes without very large study populations.

The coding variants we have identified are heterozygous in affected individuals, suggesting that either the levels of wild-type proteins are rate-limiting for somatic expansion and the variants are hypomorphic, or the variants have dominant negative or gain-of-function effects. Transcriptome-wide association studies in HD have previously implicated expression of *FAN1* and *MSH3* in modification of motor onset and progression, respectively, which supports a model of modifier gene dosage being important for somatic expansion rates (Flower et al., 2019; GeM-HD Consortium, 2019; Goold et al., 2019). Notably, the seven individuals carrying a loss-of-function variant in *MSH3* in whom wild-type MSH3 protein levels are predicted to be reduced by 50% (assuming no compensatory upregulation of the wild-type allele) all had a late/less severe phenotype, supporting previous genetic data implicating *MSH3* in HD onset and progression (Hensman Moss et al., 2017; Flower et al., 2019; GeM-HD Consortium, 2019). MSH3 forms a heterodimer with MSH2 (MutSβ) which can bind slipped DNA structures formed from trinucleotide repeats and induce expansion-prone repair (Panigrahi et al., 2010; Zhao et al., 2015). Loss-of-function variants could disrupt this process and slow or prevent the stable formation of expansion-licensed DNA substrate. Rare variants in *PMS1* were also mostly found in patients with late/less severe phenotype (Table S5), suggesting a role for PMS1 in mediating repeat expansion, as previously shown in a mouse embryonic stem cell model of Fragile X disorders (Miller et al., 2020). PMS1 forms a heterodimer with MLH1 (MutLβ), but unlike PMS2 and MLH3, the other two binding partners of MLH1, PMS1 lacks a known catalytic activity and does not seem to participate directly in mismatch repair (Räschle et al., 1999). It remains unclear how PMS1 might lead to repeat expansions.

Genetic studies have strongly implicated *FAN1* as the modifier gene at the chromosome 15 locus, with at least 4 independent GWAS signals (GeM-HD Consortium, 2019; Kim et al., 2020). Our exome-wide analysis of rare damaging variants highlighted *FAN1* as the only gene with p< 1E-3 in both dichotomous and continuous analyses (Table S7). A neighbouring gene, *TRPM1*, was also nominally significant (Table S4) but evidence from human and mouse studies strongly implicate *FAN1* as the modifier at this locus (Kim et al., 2020; Loupe et al., 2020). The overall rare, predicted damaging, variant burden in *FAN1* was three-fold higher than expected from population frequencies (gnomAD; (Karczewski et al., 2020)), further suggestive of a modifier effect. Notably, although we find the distribution of the R377W and R507H alleles heavily skewed to individuals with early onset, they are also seen in a few individuals with late onset (Figure 3), implying that the effects exerted by these variants are themselves modifiable. We observed that of the five individuals with late/less severe phenotype, R377W or R507H FAN1, and genotyping data, four had a second potential modifier that could explain the phenotype. For example, one individual carried a second CAACAG cassette after the expanded *HTT* CAG repeat. Larger studies are needed to investigate interactions between putative modifiers.

Knocking out *FAN1* in Q109 human iPSCs or iPSC-derived neurons significantly increases CAG expansion rate (Figure 5), extending the result seen in the same cell line with shRNA-mediated knockdown of *FAN1* (Goold et al., 2019). How might wild-type FAN1 slow the rate of repeat expansion? FAN1 was identified as a structure-specific DNA nuclease involved in interstrand crosslink repair and has subsequently been shown to have a role in replication fork restart and maintenance of genomic stability (MacKay et al., 2010; Smogorzewska et al., 2010; Lachaud et al., 2016). These functions have all required FAN1 nuclease activity. The clustering of rare modifier variants in FAN1 domains in our study gives new insight into how FAN1 might operate in HD. One cluster between residues 377-654 contains 11 different variants that are found in 34 individuals with early/more severe phenotype and nine individuals with late/less severe phenotype (Figure 3). Variants in this region of FAN1 might affect DNA binding or FAN1 dimerisation given that these are mediated by the SAP domain (amino acids 462-538) and a dimerisation loop (amino acids 510-518), respectively. The most common variant in this cluster, R507H, has previously been shown to reduce FAN1-DNA binding in an *in vitro* system using cell extracts (Kim et al., 2020). We showed that R507H sensitises dividing cells to mitomycin C (which induces ICLs) and also reduces nuclease activity, fitting with prior human phenotypic data suggesting that R507H FAN1 has impaired function and clinical impact (Bastarache et al., 2018).

A second cluster containing four different variants in the protein-protein interaction domain (amino acids 658-794) associated with late/less severe phenotype was observed (Figure 3). Surprisingly, these variants seem to act in the opposite direction to most damaging FAN1 variants. Given that FAN1 is known to interact with MLH1, MLH3, and PMS1 (Cannavo et al., 2007), all of which promote repeat expansions, it is possible that variants in the protein interaction domain modulate these interactions and indirectly impact on somatic expansion. Alternatively, these variants might have clustered by chance and not be true modifiers. Larger studies are required to investigate this putative signal further.

A final cluster of five different deleterious variants was observed in the C-terminal nuclease domain of FAN1 (amino acids 895-1017). These were found in five individuals with early/more severe phenotype suggesting that FAN1 nuclease activity might be required for FAN1 to slow repeat expansion and hence modify HD phenotype. We tested this hypothesis by mutating endogenous FAN1 to a nuclease-inactive form (D960A) in the Q109 iPSC model of HD. This led to a significant increase in the rate of CAG repeat expansion (Figure 5E). Notably, homozygous D960A FAN1 stimulated repeat expansion to the same extent as homozygous *FAN1* knockout, strongly suggesting that FAN1 nuclease activity is required to mediate its role in slowing repeat expansion. This notion is strengthened by the significant correlation between the nuclease activity of purified FAN1 variants and the residual age at motor onset of HD in individuals carrying those variants (Figure 4). However, previous data from a *FAN1^-/-^* osteosarcoma cell line transduced with a 118 CAG repeat showed that overexpression of wild-type, D960A or R507H FAN1 was equally effective at slowing repeat expansion (Goold et al., 2019). These conflicting findings could be reconciled if overexpression of FAN1 results in indirect effects. When FAN1 is expressed from its endogenous promoter it could be rate-limiting for repeat stability and require its nuclease activity to process DNA intermediates to prevent expansions. However, overexpression of FAN1, or its variants, could bind and sequester known interacting proteins such as PMS1, MLH1 and MLH3 (Cannavo et al., 2007) independently of FAN1 nuclease activity, either at the CAG repeat or elsewhere in the cell (Goold et al., 2019). Given that these interacting proteins normally promote repeat expansions (Pinto et al., 2013; Zhao and Usdin, 2018; Miller et al., 2020), sequestration by FAN1 or its variants might effectively inactivate them and slow expansions, as observed (Goold et al., 2019).

Our data highlight an important role for FAN1 in modulating somatic expansion in HD and show for the first time that its nuclease activity is critical for inhibition of *HTT* CAG repeat expansion. Such a role is likely to extend to other repeat expansion disorders, of which there are currently 48 known in humans (Khristich and Mirkin, 2020). For example, *FAN1* variants are associated with altered age at onset of CAG expansion-related spinocerebellar ataxias (Bettencourt et al., 2016), and FAN1 inhibits somatic expansion of repeats in mouse models of HD and Fragile X disorders (Zhao and Usdin, 2018; Loupe et al., 2020). As a structure-specific 5’-exo/endo-nuclease, FAN1 might mediate its anti-expansion effects by processing DNA repeat expansion intermediates with the flaps that are its known substrate. These intermediates could arise through repeat binding and cleavage by DNA repair factors such as MSH2/MSH3 (MutSβ) and MutL complexes (Figure 6). The resultant staggered DNA break would have repeat-containing recessed ends that could re-anneal in register (no change to repeat length) or slipped. The latter would generate a substrate for gap-filling DNA synthesis and subsequent repeat expansion, or 5’- or 3’-flaps that are known substrates for FAN1. Cleavage of these flaps by FAN1 would lead to repeat contractions and could shift the equilibrium between repeat expansion and contraction towards the latter, helping to maintain or reduce expanded repeat lengths (Figure 6). This model predicts that factors favouring repeat expansions, such as MSH3, MLH1, and an expandable *HTT* CAG repeat, would be epistatic to FAN1 in determining repeat stability as our data on non-canonical repeat sequences, and recent data from HD mouse models (Loupe et al., 2020), suggest.

**Figure 6.**
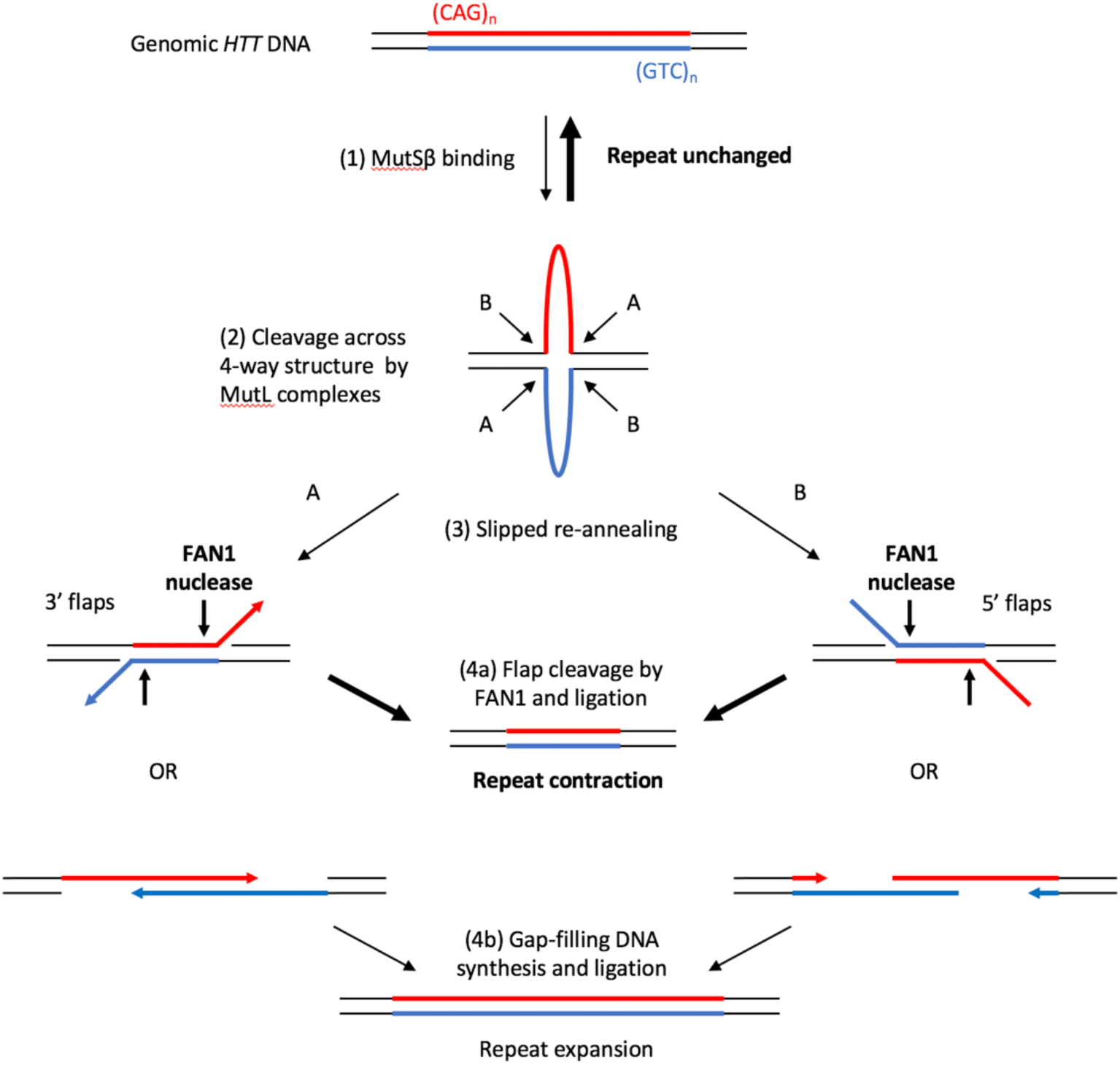
Model of how FAN1 nuclease activity could prevent CAG repeat expansions. The fully base-paired (CAG)•(CTG) tract is in dynamic equilibrium with a 4-way junction that includes loop-outs of (CAG)_n_ and (CTG)_n_ on their respective strands. Under normal cellular conditions most repeats are in their native double-stranded conformation. However, when a longer repeat tract (>35 CAG) is present it can adopt a more stable 4-way structure that can be further bound and stabilised by MSH2/MSH3 (MutSβ) (1). This 4-way junction can be cleaved on both strands in either of two orientations (A or B) by MutL complexes (2). The resulting DNA products have long overhangs, either 3’ (A) or 5’ (B), and they can either anneal fully to re-form the starting genomic DNA (top) or they can slip before partial reannealing (3). Slipped products can have 5’- or 3’-flaps and these are a substrate for FAN1 nuclease cleavage (bold arrows) and subsequent ligation, yielding repeat contractions (4a). Alternatively, the slipped products can have gaps and these are substrates for gap-filling DNA polymerases, with subsequent ligation yielding repeat expansions (4b).

In conclusion, the genetic architecture of the onset modifier trait in HD is similar to that of other oligo- or poly-genic diseases or traits, consisting of both common variants of small effect, and rarer variants of larger effect identified here (Marouli et al., 2017). The finding that single, heterozygous, coding variants in modifier genes can be associated with clinically relevant changes in HD onset or severity suggests that drugs targeting individual modifiers, or their regulators, could be effective. Such therapeutics are already in development.

## Acknowledgments

We thank all the research participants who contributed data to this research. FAN1 proteins were made by the MRC Protein Phosphorylation and Ubiquitylation Unit, University of Dundee. We acknowledge the support of the Supercomputing Wales project, which is part-funded by the European Regional Development Fund (ERDF) via Welsh Government. B.Mc. was supported by a PhD studentship from Cardiff University and an Alzheimer’s Research UK pump-priming award. J.D. was supported by a Wellcome Trust studentship. J.F.G., M.E.M. and J.D.L. received support from NIH grants NS091161 and NS082079 and from the CHDI Foundation, Inc. J-M.L. received support from grant NS105709. V.C.W. received support from grant NS049206. J.S.P. received support from NS040068 to see Predict-HD participants. A.E.R. received support from MRC, Wellcome Trust, Horizon 2020, JPND and The B.R.A.I.N. unit, funded through Health and Care Research Wales. L.J., N.M.W. and P.H. were supported by an MRC Centre grant (MR/L010305/1). M.C., D.G.M., N.D.A., L.J. and T.H.M. have all received support from CHDI. L.J. and T.H.M. received funding from the Brain Research Trust. T.H.M. was supported by a Welsh Clinical Academic Training Fellowship, an MRC Clinical Training Fellowship (MR/P001629/1) and a Patrick Berthoud Charitable Trust Fellowship through the Association of British Neurologists.

## Author Contributions

Conceptualization: M.E.M., J.D.L., E.H.A., J.F.G., N.D.A., P.H., L.J., T.H.M.

Methodology: B.Mc., J.D., C.S.B., S.P., U.C., G.E., J.S., L.E., L.-N.S., G.M., M.C., A.M., G.M., M.J.C., E.P.H., J.M.L., M.E.M., J.D.L., E.H.A., J.F.G., V.C.W., P.H., L.J., T.H.M.

Software:

Validation:

Formal Analysis: B.Mc., J.D., C.S.B., G.E., S.L., J.D.L., J.M.L., M.J.C., E.P.H., J.F.G., M.E.M., V.C.W.

Investigation: B.Mc., J.D., C.S.B., S.P., U.C., G.E., J.S., L.E., L.-N.S., M.C., A.M., D.L., M.J.C., E.P.H., T.H.M.

Resources: D.L., B.L., G.B.L., A.E.R., J.S.P., Registry and Predict investigators

Data Curation: B.Mc., J.D., J.M.L., M.J.C., E.P.H.

Writing – Original Draft Preparation, B.Mc., J.D., P.H., L.J., T.H.M.

Writing – Review & Editing: J.D., E.R., J.D.L., A.E.R., D.G.M., J.F.G., M.E.M., J.M.L., V.C.W., N.D.A.

Visualization: B.Mc., J.D., G.M.

Supervision: J.F.G., M.E.M., J.M.L., D.G.M., N.D.A., N.M.W., P.H., L.J., T.H.M.

Project Administration: J.F.G., M.E.M., J.M.L., L.J., T.H.M.

Funding Acquisition: J.F.G., M.E.M., E.H.A., J.D.L., P.H., N.D.A., N.M.W., L.J.,T.H.M.

## Declaration of Interests

V.C.W.: Scientific Advisory Board member of Triplet Therapeutics, a company developing new therapeutic approaches to address triplet repeat disorders such Huntington’s disease and Myotonic Dystrophy. Her financial interests in Triplet Therapeutics were reviewed and are managed by Massachusetts General Hospital and Mass General Brigham in accordance with their conflict of interest policies. She is a scientific advisory board member of LoQus23 Therapeutics and has provided paid consulting services to Alnylam.

J-M.L. Scientific Advisory Board of GenEdit, Inc.

J.D.L.: paid Advisory Board member for F. Hoffmann-La Roche Ltd and uniQure biopharma B.V., and he is a paid consultant for Vaccinex Inc, Wave Life Sciences USA Inc, Genentech Inc, Triplet Inc, and PTC Therapeutics Inc.

E.H.A. serves on a Data Safety Monitoring Board for Roche.

G.B.L.: Consulting services, advisory board functions, clinical trial services and/or lectures for Allergan, Alnylam, Amarin, AOP Orphan Pharmaceuticals AG, Bayer Pharma AG, CHDI Foundation, GlaxoSmithKline, Hoffmann-LaRoche, Ipsen, ISIS Pharma, Lundbeck, Neurosearch Inc, Medesis, Medivation, Medtronic, NeuraMetrix, Novartis, Pfizer, Prana Biotechnology, Sangamo/Shire, Siena Biotech, Temmler Pharma GmbH and Teva Pharmaceuticals. He has received research grant support from the CHDI Foundation, the Bundesministerium für Bildung und Forschung (BMBF), the Deutsche Forschungsgemeinschaft (DFG), the European Commission (EU-FP7, JPND). His study site Ulm has received compensation in the context of the observational Enroll-HD Study, TEVA, ISIS and Hoffmann-Roche and the Gossweiler Foundation. He receives royalties from the Oxford University Press and is employed by the State of Baden-Württemberg at the University of Ulm.

A.E.R.: Chair of European Huntington’s Disease Network (EHDN) executive committee, Global PI for Triplet Therapeutics

J.S.P. has provided consulting services, advisory board functions and clinical trial services for Acadia, Hoffman-LaRoche, Wave Life Sciences, and CHDI.

J.F.G.: Scientific Advisory Board member and has a financial interest in Triplet Therapeutics, Inc. His NIH-funded project is using genetic and genomic approaches to uncover other genes that significantly influence when diagnosable symptoms emerge and how rapidly they worsen in Huntington’s disease. The company is developing new therapeutic approaches to address triplet repeat disorders such Huntington’s disease, myotonic dystrophy and spinocerebellar ataxias. His interests were reviewed and are managed by Massachusetts General Hospital and Mass General Brigham in accordance with their conflict of interest policies. He has also been a consultant for Wave Life Sciences USA.

D.G.M.: Within the last five years D.G.M. has been a scientific consultant and/or received honoraria/stock options/research contracts from AMO Pharma, Charles River, LoQus23, Small Molecule RNA, Triplet Therapeutics and Vertex Pharmaceuticals.

L.J. is a member of the scientific advisory boards of LoQus23 Therapeutics and Triplet Therapeutics.

T.H.M. is an associate member of the scientific advisory board of LoQus23 Therapeutics.

B.Mc., J.D., C.S.B., S.P., U.C., G.E., J.S., S.L., L.E., L.-N.S., E.R., G.M., M.C., A.M., M.J.C., E.P.H., D.L., M.E.M., N.M.W., N.D.A., P.H.: nothing to disclose

## Supplemental Figures

**Figure S1.**
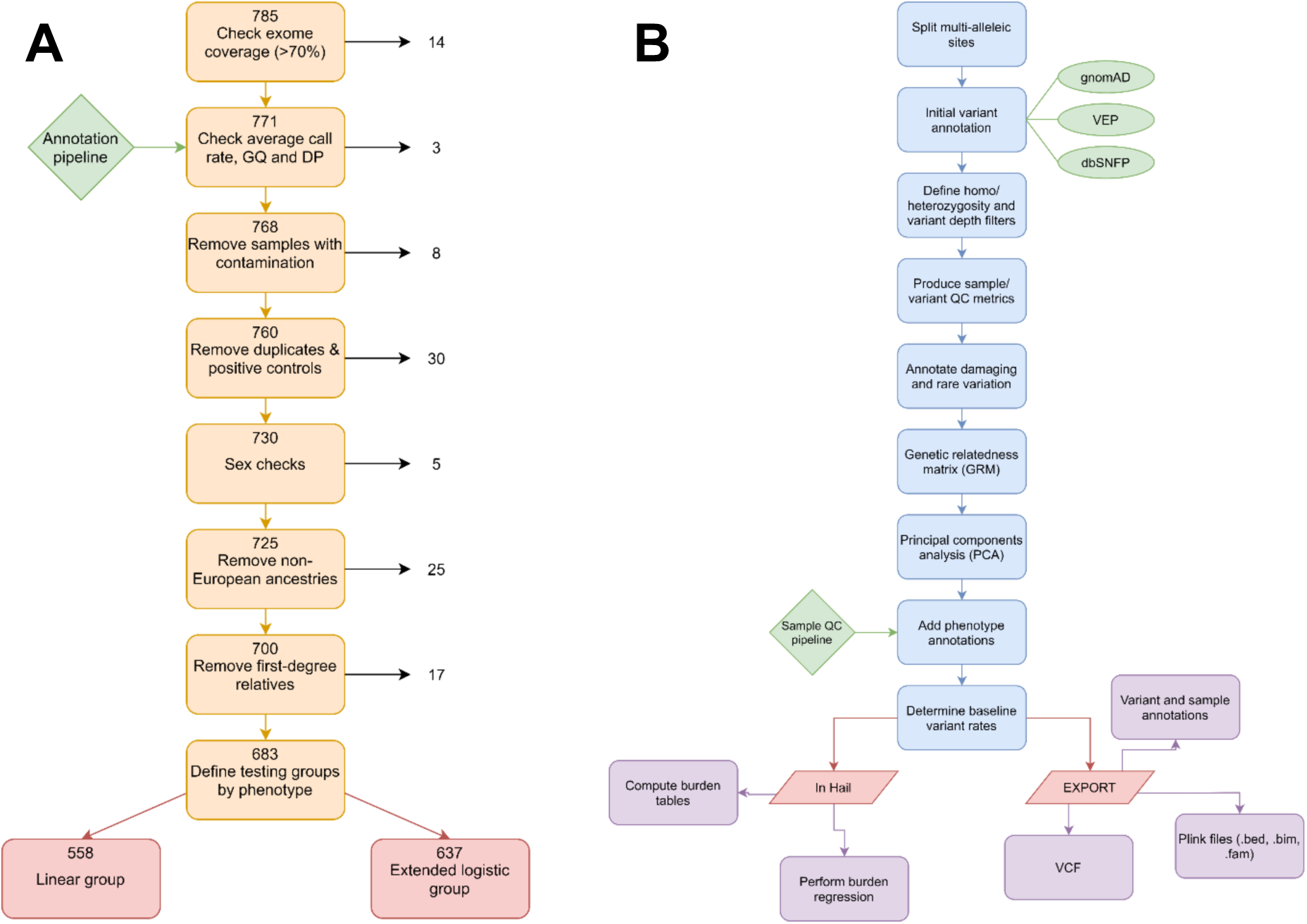
Quality control and annotation pipelines for exome sequencing data. (A) Quality control pipeline showing where and why sequencing samples were removed from the dataset. From an initial 785 sequenced exomes, 683 passed all quality control steps. Subgroups of this population were used in downstream analyses: a continuous group (N=558) containing all individuals with a known age at motor onset and a dichotomous group (N=637) containing all individuals with an extreme phenotype, either early or late actual or predicted onset of symptoms, or more or less severe motor or cognitive symptom scores. (B) Annotation pipeline indicating the pathway, databases (gnomAD & dbSNFP) and tools (variant effect predictor tool (VEP)) used to annotate individual variants across exomes.

**Figure S2.**
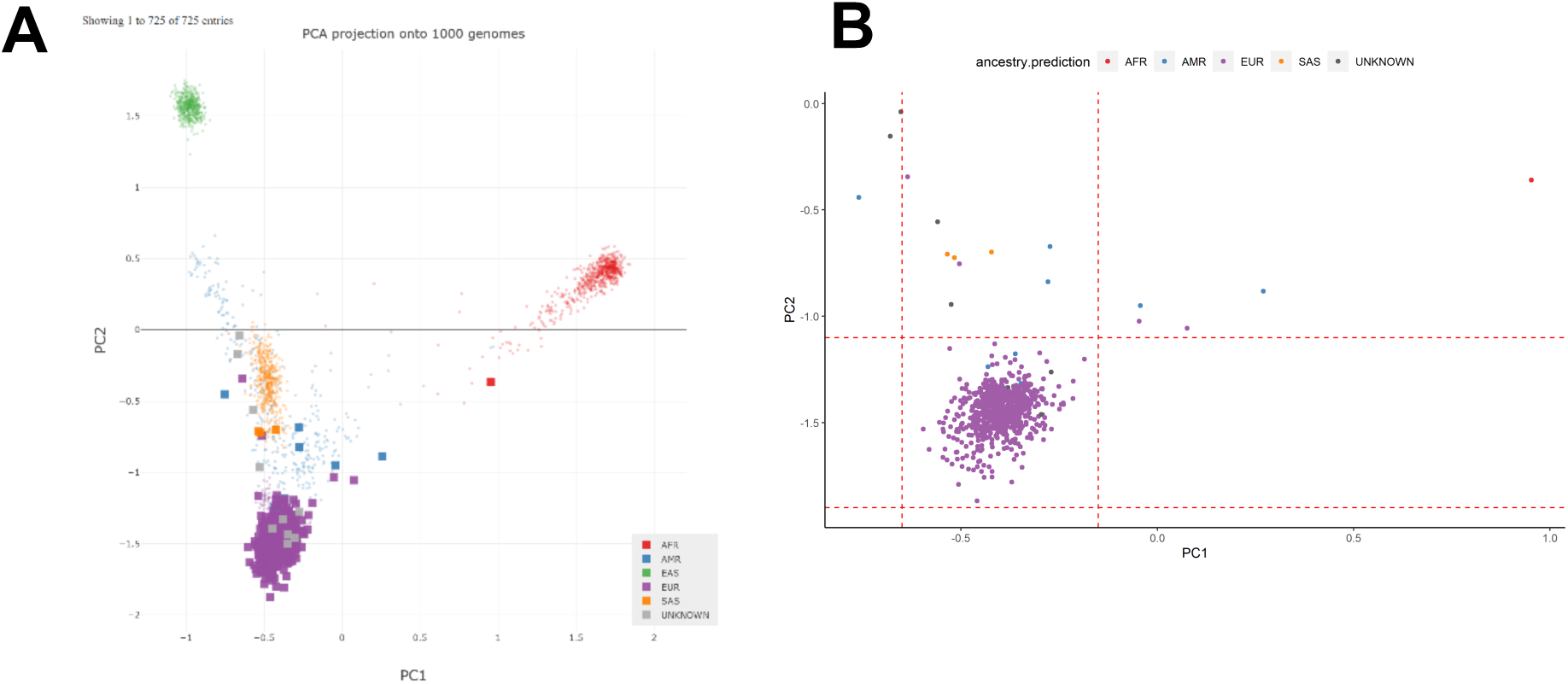
Individuals of European Ancestry were selected for downstream analysis. (A) Principal component analysis and estimated ancestries against the 1,000 genome project as calculated by Peddy(Pedersen and Quinlan, 2017) for N=725 individuals passing initial quality control (See Figure S2A). Individuals of European ancestry in purple. (B) Magnified area of the principal component analysis (PC1 and PC2) from (A). Individuals with predicted European ancestry enclosed within the rectangle bordered by dotted lines were retained in analyses (N=700). Key: AFR, African; AMR, Ad Mixed American; EAS, East Asian; EUR, European; SAS, South Asian

**Figure S3.**
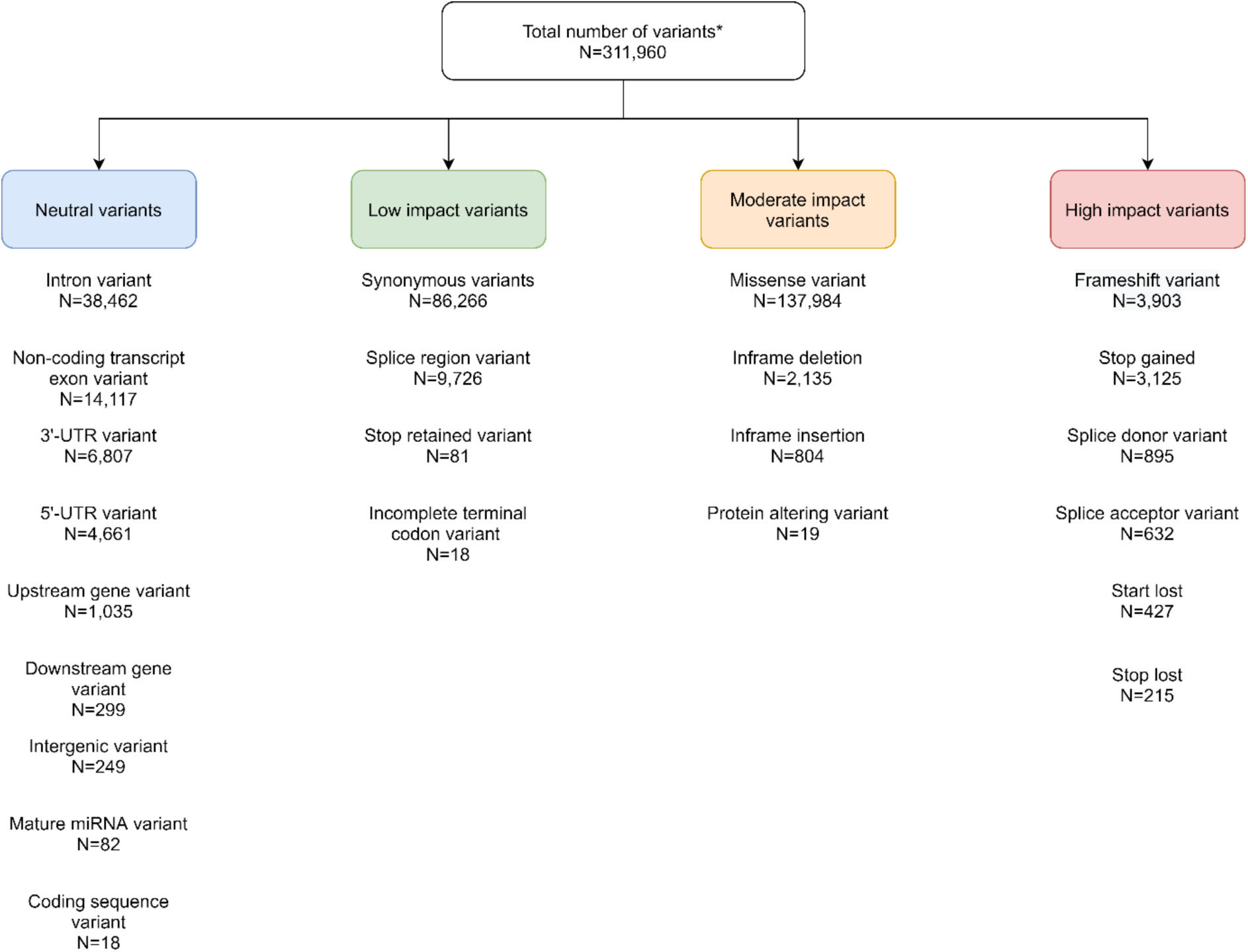
The types and predicted functional impacts of all variants identified through exome sequencing. All variants compared with the hg19 human genome were annotated using the pipeline in Figure S1B and divided into four classes based on predicted functional impact. For variants with multiple annotations, the most damaging annotation (listed first in variant effect predictor (VEP)) was used. Total variant numbers are shown for the 683 exomes which passed quality control.

**Figure S4.**
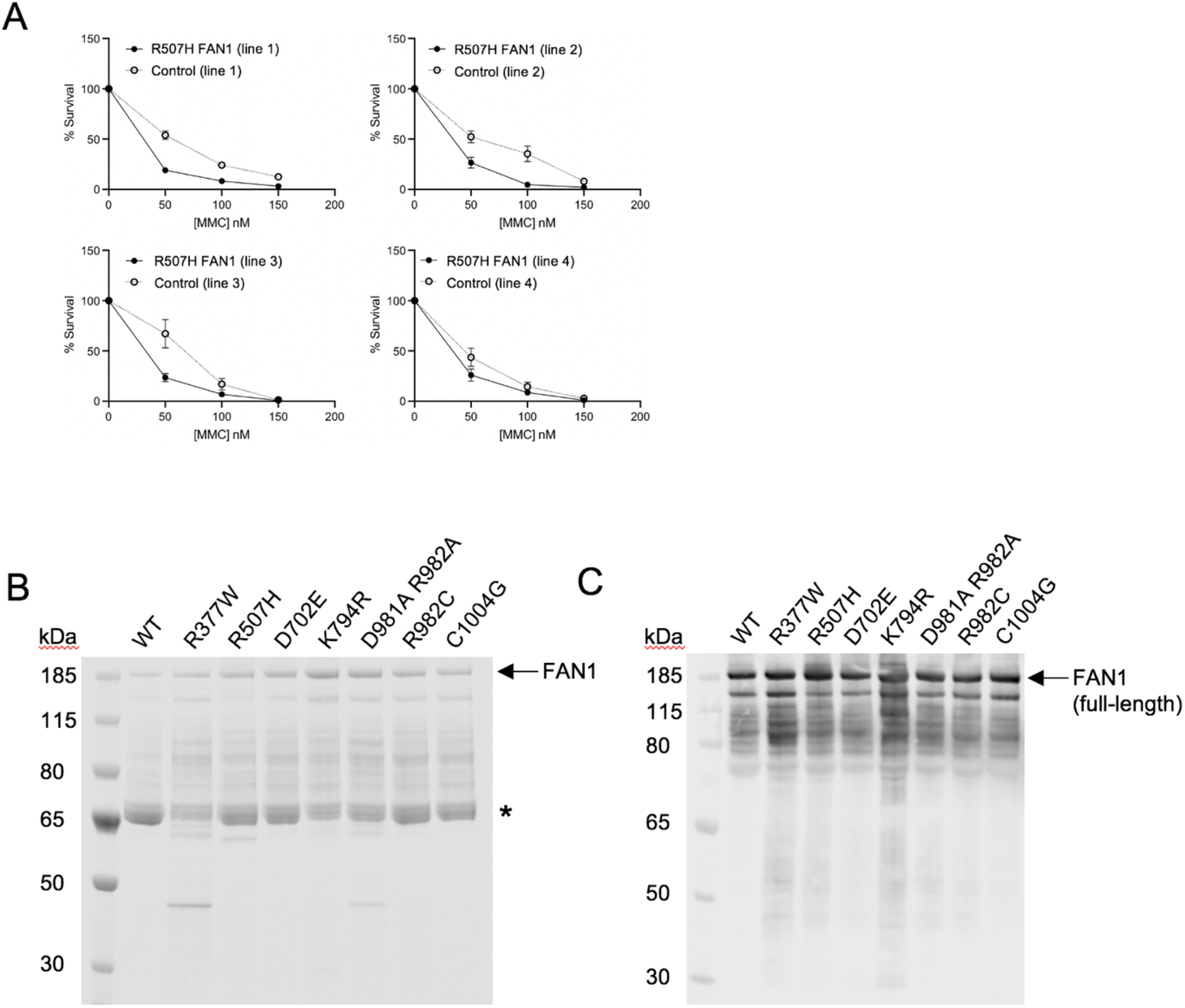
HD lymphoblastoid cells carrying R507H are sensitive to mitomycin C. Partial purification of wild-type and FAN1 variants. (A) 4 lymphoblastoid cell lines derived from individuals with HD carrying a heterozygous R507H FAN1 variant were grown alongside control HD lines with homozygous wild-type FAN1 matched for age and pure CAG length and treated with a single dose of 0-150 nM mitomycin C. Survival (% of untreated live cell count) at 7 days shown for each line. (B) NusA-His-tagged full-length FAN1 proteins (wild-type (WT) and variants as shown) were expressed in *E. coli* and partially purified using cobalt-agarose. Samples were run on 4-12% Bis-Tris SDS-PAGE and stained with Coomassie blue. Expected size of NusA-His-FAN1 is 175.9 kDa as indicated. Main contaminants (*) identified by mass spectrometry as *E. coli* chaperones ArnA and DnaK. Further purification of FAN1 proteins was attempted but active protein yields were low. (C) Immunoblot of WT and FAN1 variants run on 4-12% Bis-Tris SDS-PAGE as in (B). Primary antibody was sheep polyclonal to FAN1 (CHDI). Full-length FAN1 is seen along with multiple degradation products.

**Figure S5.**
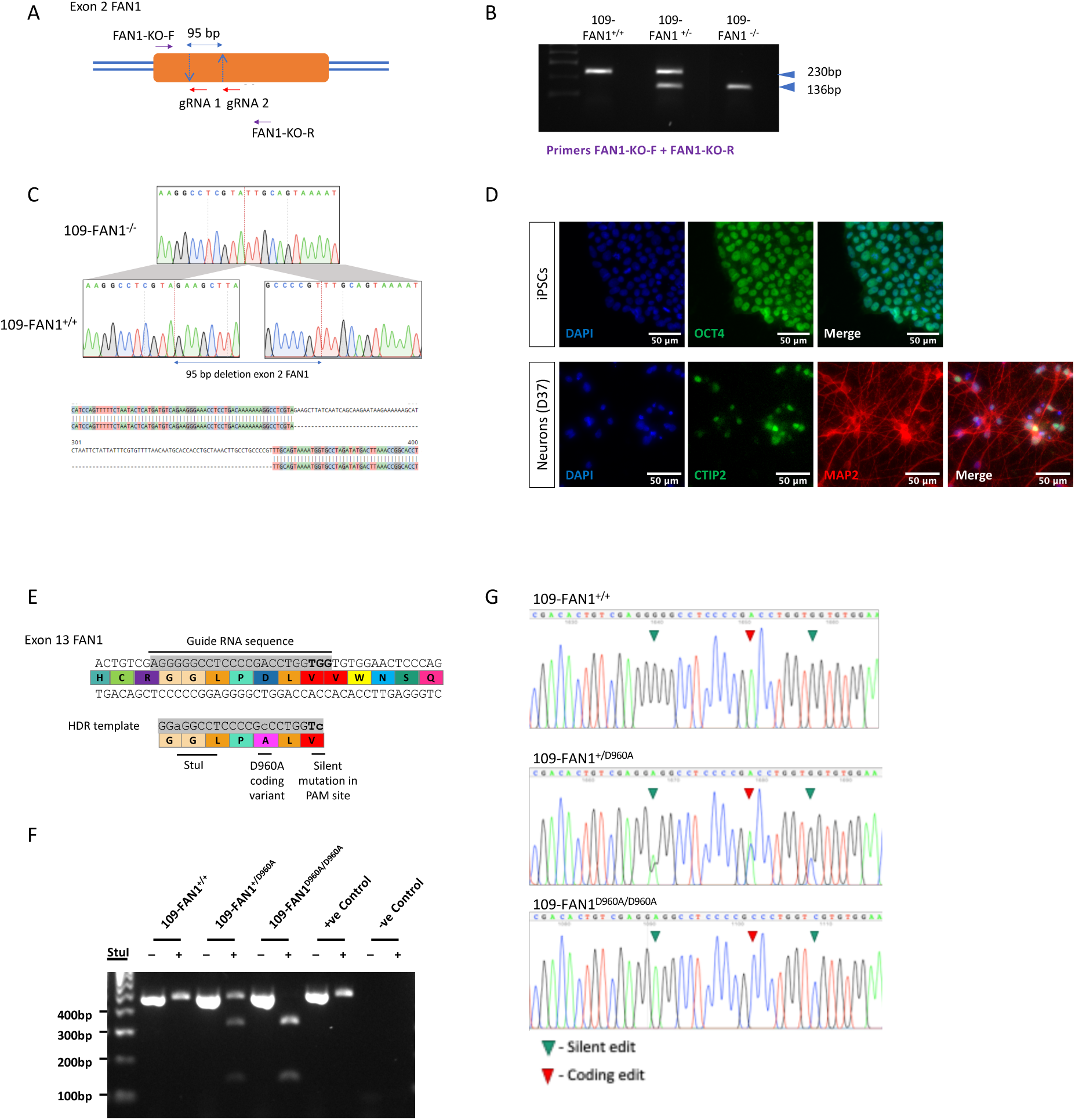
Knockout and editing of *FAN1* in Q109 HD iPSC line using CRISPR/Cas9. (A) Schematic depicting CRISPR-Cas9 targeting of exon 2 of *FAN1* using two guide RNAs (gRNAs) and the primer pair used to amplify exon 2 of *FAN1*. (B) Diagnostic PCR screen using primers, FAN1-KO-F + FAN1-KO-R highlights wild-type (WT), *FAN1*^-/-^ and *FAN1*^+/+^ candidate clones. Representative banding patterns of WT alleles (230bp), *FAN1*^+/-^ (230/136 bp) and *FAN1*^-/-^ (136/136 bp). (C) Sanger sequencing of PCR products demonstrates the targeted 95 bp deletion in exon 2 of *FAN1*. (D) Q109 iPSCs are pluripotent whereas differentiated neurons (day 37) display neuronal markers. Undifferentiated Q109 iPSCs stained for the pluripotency marker OCT4 (top row). iPSC-derived neurons stained for the neuronal marker MAP2 (red) and CTIP2 (green). All nuclei are counterstained with DAPI (blue). (E) Construction of FAN1 D960A edit. CRISPR/Cas9 was used to target exon 13 of *FAN1* using a homology directed repair (HDR) template. A single guide RNA sequence (grey) and a 122 bp HDR template containing the desired gene edit coding for a D960A point mutation were utilised to generate edited Q109 cells. The HDR template contained two silent mutations (lowercase) to prevent Cas9 re-cutting of the edited region and to introduce a *StuI* restriction digest site for diagnostic screening. (F) Restriction digest with *StuI* confirms 109-FAN1^+/+^, 109-FAN1^+/D960A^ and 109-FAN1^D960A/D960A^ genotypes. StuI cleaves the 442bp PCR product into 124 and 318bp products only in the presence of the silent 2868G>A mutation. (G) Sanger sequencing of PCR products confirms introduction of coding change for D960A variant.

## Supplemental Tables

**Table S1.** CAG repeat sequence retains a significant association with age at HD onset after accounting for pure CAG length. Age at motor onset was regressed on polyglutamine (PolyQ) or pure CAG (PolyCAG) length with or without covariates to represent non-canonical allele sequences. Covariate I_0 indicates the presence (1) or absence (0) of an expanded *HTT* allele lacking a CAACAG cassette (Figure 2, allele groups c-d). Covariate I_2 indicates the presence (1) or absence (0) of an expanded *HTT* allele with additional 3’ non-canonical CAACAG, or CAA or CAC trinucleotides (see also Figure 2, allele groups e-h). All samples passing quality control that had been through ultra-high MiSeq depth sequencing or had repeat lengths and sequences confidently called from PCR-fragment length analysis and exome sequencing data were included (Total N=558; Registry-HD N=463, Predict-HD N=95). Significant p-values after multiple testing correction for 5 models (p <1E-02) in bold.

| Model | Covariate | B | p | R<sup>2</sup> |
| --- | --- | --- | --- | --- |
| Ln(AMO) ~ PolyQ length | PolyQ | -0.083 | **7.15E-47** | 0.310 |
| Ln(AMO) ~ PolyCAG length | CAG | -0.092 | **3.09E-58** | 0.372 |
| Ln(AMO) ~ PolyCAG length + I_0 | CAG | -0.091 | **1.90E-57** | 0.382 |
|  | I_0 | -0.177 | **4.04E-03** |  |
| Ln(AMO) ~ PolyCAG length + I_2 | CAG | -0.092 | **2.67E-59** | 0.386 |
|  | I_2 | 0.233 | **2.04E-04** |  |
| Ln(AMO) ~ PolyCAG length + I_0 + I_2 | CAG | -0.091 | **1.68E-58** | 0.393 |
|  | I_0 | -0.168 | **5.93E-03** |  |
|  | I_2 | 0.225 | **2.97E-04** |  |

**Table S2.** Demographic details for individuals passing quality control and included in the exome sequencing analyses. Total N = 683 (465 Registry-HD group; 218 Predict-HD group). Note that not all individuals were part of the extreme phenotype groups (early/late in Registry-HD, more severe/less severe in Predict-HD) after correction for pure CAG repeat lengths determined by MiSeq or exome sequencing. Mean (standard deviation) shown for pure CAG length after sequencing, age at motor onset (AMO) and residual AMO. Residual AMO is calculated by subtracting age at onset predicted by CAG length alone from actual age at onset, determined clinically.

| Study Group | N | Sex (% male)<sup>a</sup> | CAG length | Age at Motor Onset (AMO)<sup>b</sup> | Residual AMO<sup>b</sup> |
| --- | --- | --- | --- | --- | --- |
| Registry-HD | 465 | 46.6% | 42.9 (2.2) | 48.4 (16.4) | -0.8 (13.4) |
| Early (Registry-HD) | 214 | 45.3% | 43.4 (2.3) | 33.7 (7.8) | -13.9 (4.9) |
| Late (Registry-HD) | 207 | 48.3% | 42.3 (1.8) | 63.9 (7.7) | +12.7 (4.2) |
| Predict-HD | 218 | 38.1% | 43.3 (2.8) | 47.1 (9.8) | -1.0 (5.9) |
| More severe (Predict-HD) | 101 | 31.7% | 43.2 (2.7) | 46.1 (9.5) | -1.7 (5.5) |
| Less severe (Predict-HD) | 115 | 42.6% | 43.4 (3.0) | 53.6 (10.1) | +5.0 (5.9) |

(a) One individual in the late Registry-HD group was of unknown sex and not included in the sex column
(b) AMO and residual AMO were calculable for all in the Registry-HD group. For the Predict-HD group, 93 had had onset and hence calculable AMO and residual AMO (80 in the more severe group; 11 in the less severe group; 2 in neither extreme group)

**Table S3.** Glutamine-encoding *HTT* repeat tract sequences for Predict-HD group. Whole-exome sequencing data were extracted and manually assessed to determine the sequence of the glutamine-encoding repeat tract in expanded *HTT* alleles of Predict-HD participants (N=216). It was not possible to determine full allelic structures as read lengths were 75 bp. Phase could be determined for atypical alleles as wild-type CAG length was short enough to be effectively captured. The presence or absence of extra non-canonical CAACAG or CAA or CAC triplets was used as a covariate in analyses (see Table S1).

| CAG tract sequence<br>(expanded *HTT*) | Predicted early or worst<br>TMS/SDMT (more severe) | Predicted late or best<br>TMS/SDMT (less severe) |
| --- | --- | --- |
| Canonical (CAACAG) | 98 | 111 |
| (CAACAG)<sub>2</sub> | 1 | 2 |
| CAC(CAG)<sub>3</sub>CAACAG | 0 | 2 |
| No interruption (pure CAG) | 2 | 0 |

**Table S4.** Candidate gene analysis for all genes found at modifier loci in GeM-HD GWAS. Optimal sequence kernel association tests (SKAT-O) were performed using rare (MAF < 0.01) for candidate modifier genes from GeM-HD GWAS. Gene-wide variant numbers were regressed on either dichotomous early/more severe or late/less severe phenotypes (logistic group, N=637) or residual age at motor onset (linear group, N=558). Phenotypes were corrected for non-canonical CAG repeats, either by using a covariate in logistic analyses, or pure CAG lengths in continuous analyses. Three different variant groups were tested: all non-synonymous (NS), non-synonymous and predicted damaging to protein function (NSD20; CADD PHRED score ≥ 20), and loss-of-function (LoF). Genes with nominal significance (p < 0.05) are in bold. Several genes from GWAS loci were not annotated in exome sequencing data and hence not included: *OSGEPL1-AS1* (2A); *JRKL-AS1* + *MIR1260B* (11A); *LOC102723562* (12A); *HERC2P10* + *LOC100288637* + *MIR211* (15A) + *MIR3591* (18A). See also Table 1.

| Chr | Gene | Dichotomous group (N=637) |  |  | Linear group (N=558) |  |  |
| --- | --- | --- | --- | --- | --- | --- | --- |
|  |  | NS | NSD20 | LoF | NS | NSD20 | LoF |
| 2A | *ANKAR* | 8.47E-01 | 8.28E-01 | NA | 8.44E-01 | 6.91E-01 | 7.86E-01 |
| 2A | *ASNSD1* | 9.01E-01 | 8.20E-01 | NA | 5.61E-01 | 6.97E-01 | NA |
| 2A | *ORMDL1* | 3.77E-01 | 4.34E-01 | NA | 4.81E-01 | 5.29E-01 | NA |
| 2A | *OSGEPL1* | 8.16E-01 | 8.28E-01 | 7.80E-01 | 7.79E-01 | 7.81E-01 | 8.47E-01 |
| 2A | *PMS1* | **2.72E-02** | **2.65E-03** | 3.13E-01 | 2.62E-01 | 1.08E-01 | NA |
| 3A | *EPM2AIP1* | 4.87E-01 | 4.51E-01 | NA | 1.00E+00 | 1.00E+00 | NA |
| 3A | *GOLGA4* | 8.36E-01 | 6.72E-01 | **3.40E-03** | 6.59E-01 | 2.18E-01 | 6.30E-02 |
| 3A | *LRRFIP2* | 3.57E-01 | 4.91E-01 | NA | 3.62E-01 | 8.85E-01 | NA |
| 3A | *MLH1* | 1.72E-01 | 5.58E-02 | NA | 4.99E-01 | 7.45E-01 | NA |
| 3A | *TRANK1* | 4.10E-01 | 5.42E-01 | 8.98E-01 | 1.97E-01 | 3.63E-01 | 6.07E-01 |
| 3A | *C3orf35* | 3.21E-01 | 2.17E-01 | 2.16E-01 | 2.01E-01 | 1.63E-01 | 1.35E-01 |
| 3A | *ITGA9* | 1.00E+00 | 8.51E-01 | NA | 8.76E-01 | 8.99E-01 | NA |
| 5A | *ANKRD34B* | 1.80E-01 | 8.61E-01 | NA | 1.64E-01 | 6.76E-01 | NA |
| 5A | *DHFR* | 1.05E-01 | 9.83E-02 | NA | 1.54E-01 | 1.42E-01 | NA |
| 5A | *FAM151B* | 1.51E-01 | 1.45E-01 | NA | 5.79E-01 | 5.77E-01 | NA |
| 5A | *MSH3* | 5.96E-02 | 2.78E-01 | **9.51E-03** | 3.00E-01 | 6.35E-01 | 1.09E-01 |
| 5A | *MTRNR2L2* | 2.22E-01 | NA | NA | 8.12E-01 | NA | NA |
| 5B | *GPR151* | 5.48E-01 | 4.20E-01 | 5.12E-01 | 2.65E-01 | 2.54E-01 | 2.12E-01 |
| 5B | *PPP2R2B* | 1.61E-01 | 2.52E-01 | NA | 4.82E-01 | 7.36E-01 | 8.41E-01 |
| 5B | *TCERG1* | **1.25E-02** | 4.54E-01 | NA | **2.74E-03** | 2.76E-01 | NA |
| 7A | *AIMP2* | 5.16E-02 | 1.75E-01 | 7.17E-01 | 1.66E-01 | 2.08E-01 | 1.58E-01 |
| 7A | *ANKRD61* | 5.25E-01 | 1.00E+00 | 2.47E-01 | 5.05E-01 | 7.08E-01 | 6.44E-01 |
| 7A | *CCZ1* | 1.55E-01 | 1.84E-01 | NA | 2.80E-01 | 2.88E-01 | NA |
| 7A | *CYTH3* | 8.43E-02 | 7.99E-02 | NA | 1.01E-01 | 1.19E-01 | NA |
| 7A | *EIF2AK1* | 6.63E-01 | 7.28E-01 | NA | 7.05E-01 | 7.31E-01 | NA |
| 7A | *OCM* | 8.61E-01 | 8.65E-01 | 1.00E+00 | 1.00E+00 | 1.00E+00 | 8.54E-01 |
| 7A | *PMS2* | 1.00E+00 | 9.00E-01 | 5.55E-01 | 5.59E-01 | 3.51E-01 | 1.33E-01 |
| 7A | *RSPH10B* | NA | NA | NA | NA | NA | NA |
| 7A | *RSPH10B2* | NA | NA | NA | 8.92E-01 | NA | NA |
| 7A | *USP42* | 7.18E-01 | 2.49E-01 | NA | 1.00E+00 | 3.89E-01 | NA |
| 8A | *RRM2B* | 1.69E-01 | NA | NA | **2.26E-02** | NA | NA |
| 8A | *UBR5* | 7.47E-01 | 7.23E-01 | NA | 4.11E-01 | 3.88E-01 | NA |
| 11A | *CCDC82* | 5.27E-01 | NA | NA | 6.09E-01 | 9.01E-01 | NA |
| 11A | *JRKL* | 2.48E-01 | 2.42E-01 | NA | 7.74E-02 | 7.07E-02 | NA |
| 11A | *MAML2* | 1.00E+00 | 8.94E-01 | 2.08E-01 | 5.16E-01 | 4.62E-01 | 1.31E-01 |
| 11B | *SYT9* | 1.05E-01 | 1.13E-01 | NA | 4.20E-01 | 4.12E-01 | NA |
| 12A | *CORO1C* | 8.75E-01 | 8.69E-01 | 8.33E-01 | 6.78E-01 | 6.66E-01 | 8.44E-01 |
| 12A | *FICD* | 3.41E-01 | 3.42E-01 | NA | 5.86E-01 | 7.55E-01 | NA |
| 12A | *ISCU* | NA | NA | NA | NA | NA | NA |
| 12A | *SART3* | 9.02E-01 | 7.23E-01 | NA | 8.15E-01 | 4.32E-01 | NA |
| 12A | *SELPLG* | 5.31E-01 | 8.55E-01 | NA | 7.91E-01 | 6.61E-01 | NA |
| 12A | *SSH1* | 6.77E-02 | 1.45E-01 | 5.09E-01 | 2.77E-01 | 4.13E-01 | 9.81E-01 |
| 12A | *TMEM119* | 5.56E-01 | NA | NA | 5.28E-01 | NA | NA |
| 15A | *FAN1* | **2.32E-04** | **6.61E-04** | 2.17E-01 | **1.09E-03** | **9.06E-04** | 3.40E-01 |
| 15A | *MTMR10* | 8.11E-01 | 7.95E-01 | NA | 3.01E-01 | 2.77E-01 | NA |
| 15A | *TRPM1* | **9.11E-03** | **3.24E-03** | 2.28E-01 | **1.41E-02** | **6.90E-03** | 3.09E-01 |
| 16A | *GSG1L* | 9.05E-01 | 9.13E-01 | NA | 4.09E-01 | 4.35E-01 | NA |
| 18A | *ALPK2* | 6.47E-01 | 1.86E-01 | 4.06E-01 | 4.74E-01 | 8.83E-01 | 8.50E-01 |
| 18A | *MIR122* | NA | NA | NA | NA | NA | NA |
| 19A | *C19orf68* | 5.15E-02 | 5.41E-02 | NA | 1.80E-01 | 1.66E-01 | NA |
| 19A | *LIG1* | 9.47E-02 | 7.07E-02 | 5.09E-01 | **3.63E-02** | **2.30E-02** | 5.16E-01 |
| 19A | *PLA2G4C* | 8.02E-01 | 8.68E-01 | NA | 3.99E-01 | 7.39E-01 | NA |

**Table S5.**
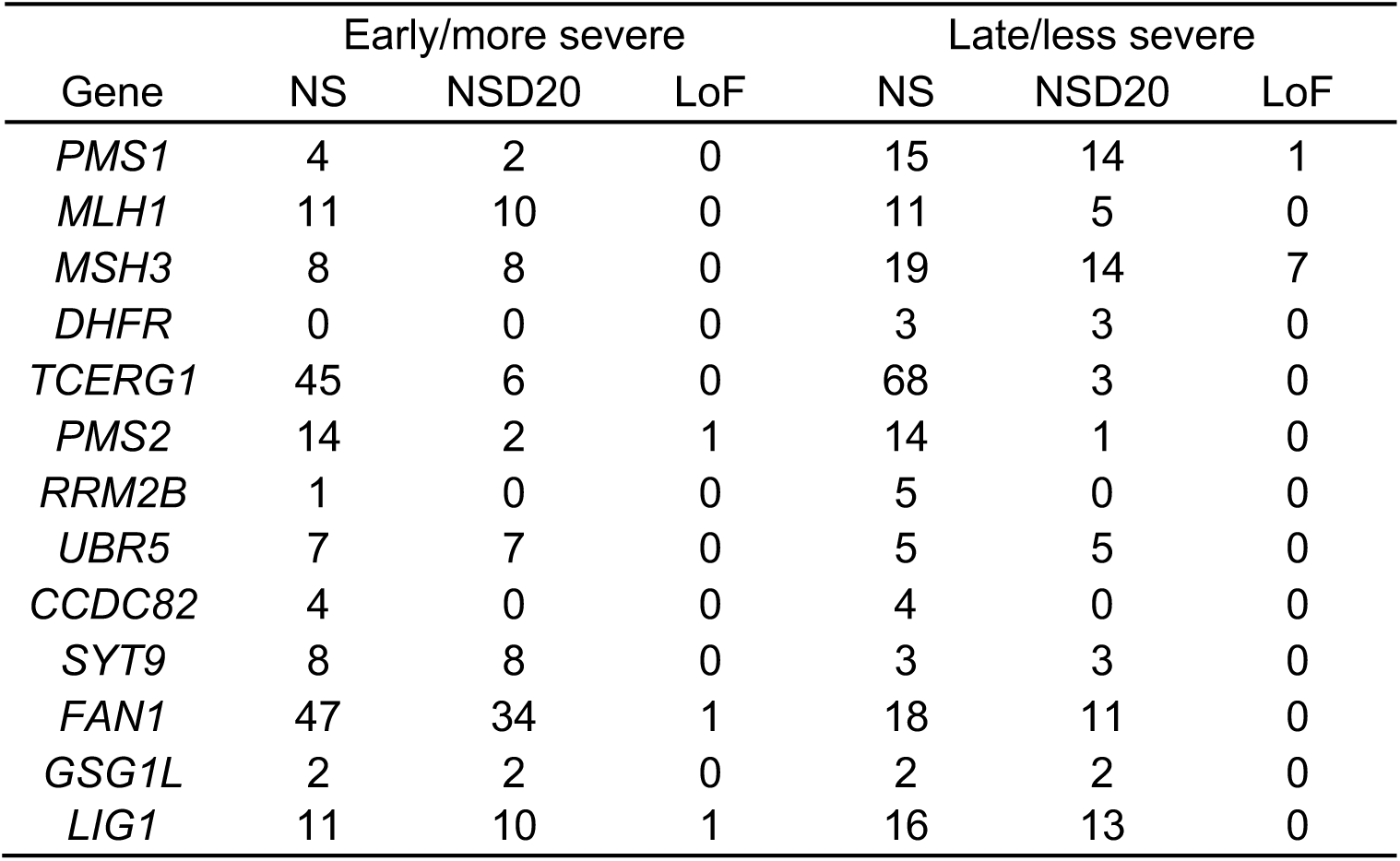
Burden of rare non-synonymous variants of different types in candidate genes in the early/more severe and late/less severe phenotype groups. The numbers of non-synonymous (NS), non-synonymous damaging (NSD20; variants either CADD PHRED score > 20 or predicted loss-of-function) and predicted loss-of-function (LoF) variants in candidate genes from significant loci in GeM-GWAS. To be included, variants had to have minor allele frequency < 0.01 and call rate >75%, as tested in SKAT-O analyses. See also Table 1.

**Table S6.**
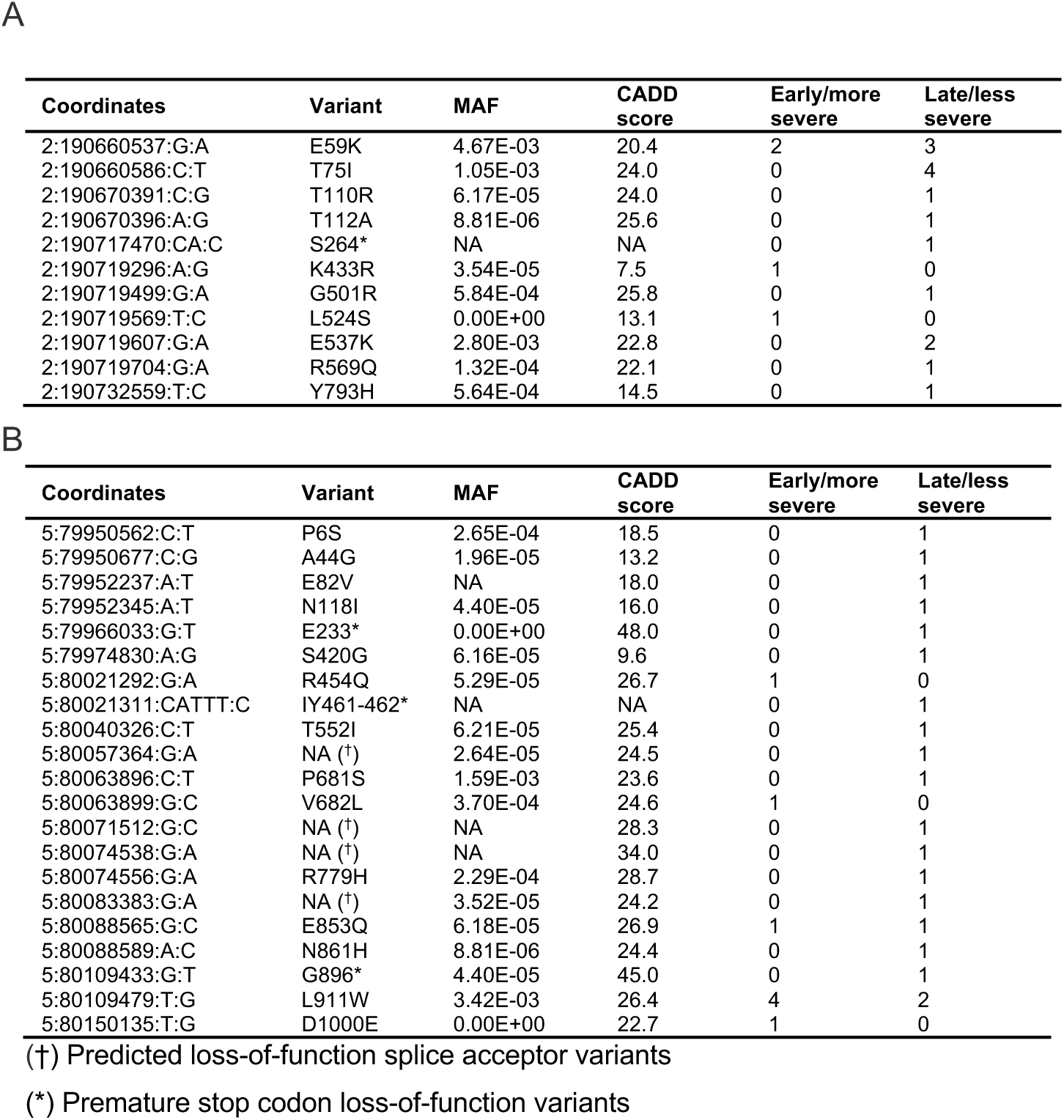
Rare non-synonymous coding variants in PMS1 and MSH3 identified in extreme phenotype groups of individuals with HD. Allele counts for all rare (MAF < 0.01) non-synonymous coding variants in (A) PMS1 and (B) MSH3 identified through exome sequencing in the dichotomous cohort (N=637). Genomic coordinates are given for hg19. Early/more severe corresponds to early or predicted early onset or high TMS or SDMT. Late/less severe corresponds to late or predicted late onset or low TMS or SDMT. CADD score predicts how damaging individual variants are to protein function. CADD score > 20 implies a variant is in the top 1% predicted most damaging substitutions in the human genome (dbSNFP v4.0(Liu et al., 2011, 2016)). Minor allele frequencies (MAF) taken from the European arm of gnomAD (Karczewski et al., 2020) v2.1.1.

**Table S7.** Top 15 genes in exome-wide analysis of association between rare predicted damaging variants and HD clinical phenotypes, corrected for CAG repeat sequence differences. Exome-wide optimal sequence kernel association tests (SKAT-O) were performed using rare (MAF < 0.01), predicted damaging, non-synonymous variants (loss-of-function and/or CADD PHRED ≥20) collapsed on genes, and testing for association with clinical phenotypes. In the dichotomous analysis, association of phenotype (early/more severe or late/less severe) was tested using logistic regression with a covariate for *HTT* CAG repeat structure to account for non-canonical CAG repeats. In the continuous analysis, linear regression was used to test association of variants with residual age at motor onset, corrected for pure CAG lengths. Covariates used: principal components (PCA) 1-5; baseline variant rate (BVR); mean variant depth; study ID (Registry or Predict). Significance threshold = 1.3E-5 (Bonferroni correction for 3912 genes with at least 10 variants).

| Dichotomous (N=637) |  | Continuous (N=558) |  |
| --- | --- | --- | --- |
| Gene | p | Gene | p |
| *DMGDH* | 2.92E-04 | *CUBN* | 1.44E-04 |
| *OR4C15* | 4.20E-04 | *ERAP2* | 3.96E-04 |
| *FAN1* | 6.61E-04 | *ZNF462* | 6.18E-04 |
| *IGF1R* | 1.07E-03 | *DENND4B* | 6.60E-04 |
| *MUC6* | 1.55E-03 | *KIAA0319* | 8.79E-04 |
| *ELP2* | 1.87E-03 | *FAN1* | 9.06E-04 |
| *LAD1* | 2.24E-03 | *FBP2* | 1.06E-03 |
| *PEG3* | 2.34E-03 | *SIPA1L2* | 1.15E-03 |
| *PMS1* | 2.65E-03 | *C9* | 1.17E-03 |
| *NDOR1* | 2.67E-03 | *NAALAD2* | 1.39E-03 |
| *C2CD3* | 3.10E-03 | *SAMD3* | 1.83E-03 |
| *FBP2* | 3.24E-03 | *MUT* | 2.10E-03 |
| *TRPM1* | 3.24E-03 | *P2RY13* | 2.32E-03 |
| *RAD54L* | 3.35E-03 | *ANXA11* | 2.78E-03 |
| *ST7L* | 3.65E-03 | *PPP2R1B* | 3.22E-03 |

**Table S8.**
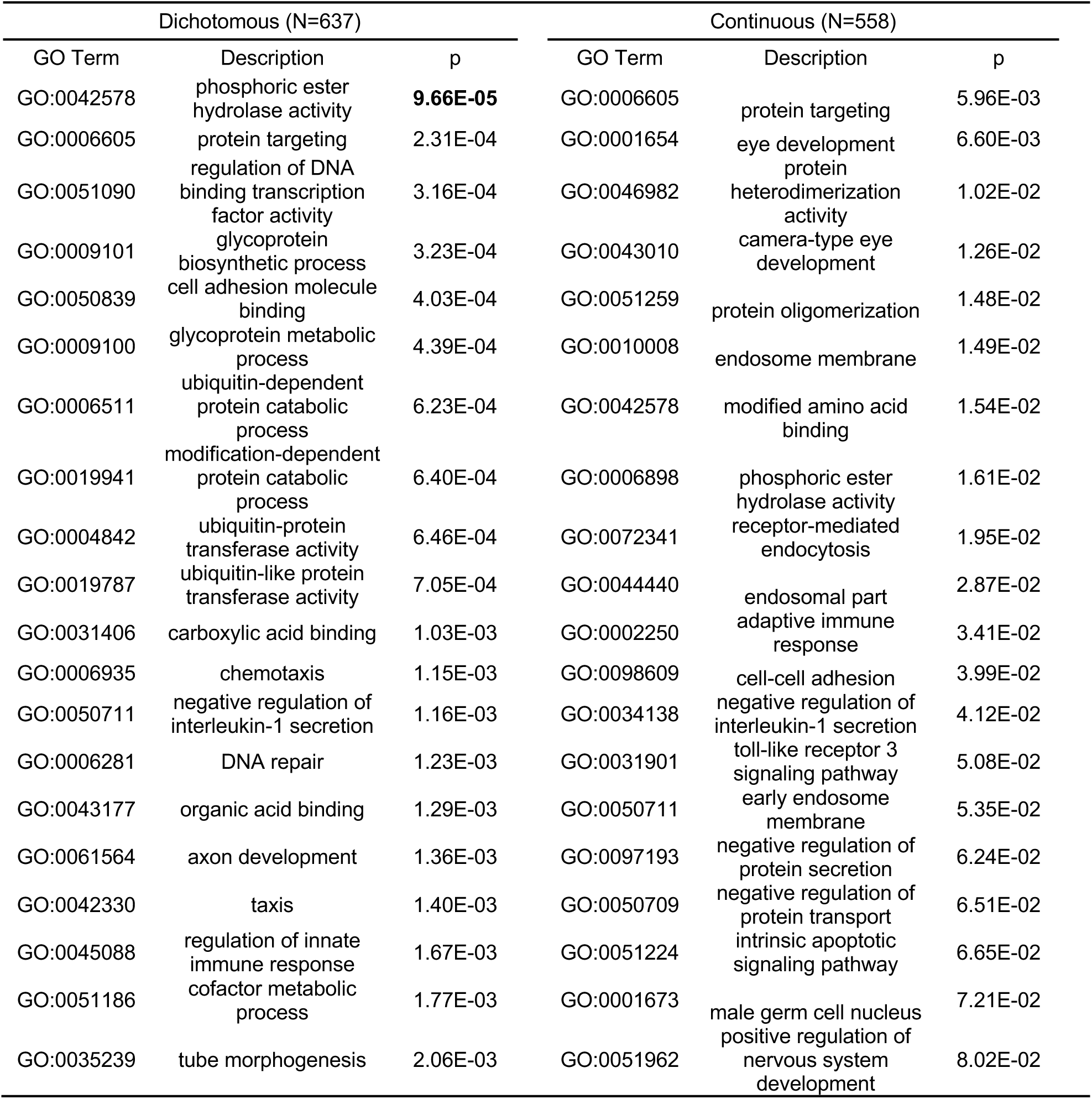
Candidate Pathway Analysis. Significant gene pathways from GWAS (GeM-HD Consortium, 2019) (*q*<1E-03) were selected as candidate pathways, and gene set membership was taken from the gene ontology (GO) database (Ashburner et al., 2000; The Gene Ontology Consortium, 2019). p values from whole-exome SKAT-O analyses (see also Table S7) were combined across GO pathways using Fisher’s method. Missing genes or genes with missing p values were excluded. Significance pathways in bold (p< 1.1E-04 (multiple testing correction for 233 pathways and 2 tests).

**Table S9.**
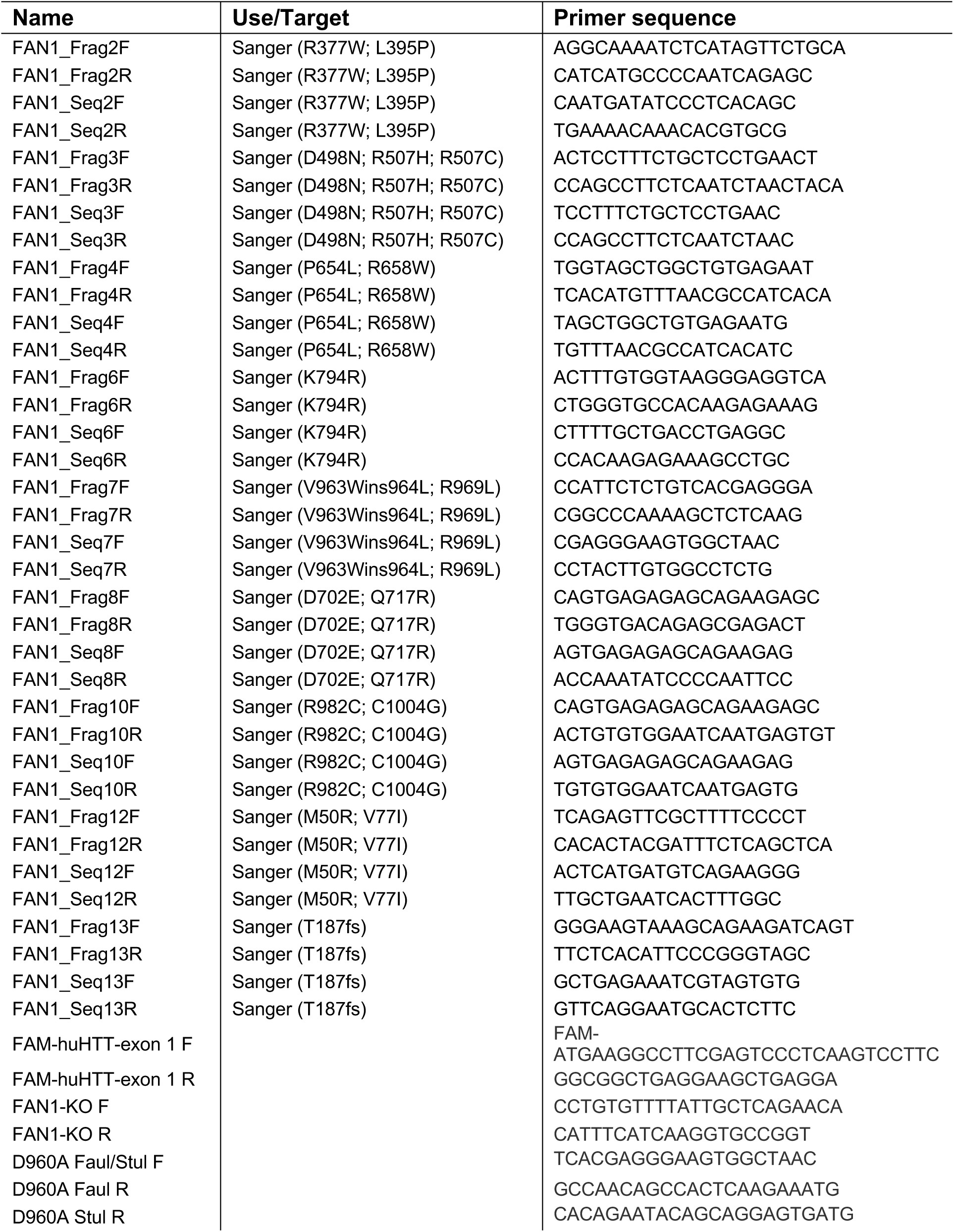
Primers used in this study. Primers marked by ‘Sanger’ were used for sanger sequencing of *FAN1* variants in select individuals. *FAN1* Q717R was confirmed both by exome QC and Sanger sequencing to not exist (this was originally called but later refuted in the QC pipeline).

## Lead contact and materials availability

Questions or requests for data or resources should be directed to the Lead Contact, Dr Thomas H. Massey. Data from human subjects will be shared with qualified researchers once appropriate assurances are made that no attempt to identify patients from which the data were derived will be made.

## Methods

### Experimental model and subject details

#### Clinical sample selection

Human subjects were selected from two HD patient cohorts. The first cohort was from the European Registry-HD study (Orth et al., 2010), which primarily enrolled individuals with HD and clinical onset. A total of 6086 individuals had both a known inherited pure CAG length (40-55 CAGs; after sequencing in this project the range was 38-55 CAGs) and an age at first onset of motor symptoms. Of these, 3046 had CAG lengths determined by BioRep in line with Registry protocols (https://www.enroll-hd.org/enrollhd_documents/2016-10-R1/registry-protocol-3.0.pdf) and 3040 had CAG lengths determined by local labs. CAG lengths on which samples were selected were determined by a standard PCR fragment length assay and assumed a canonical glutamine-encoding repeat sequence in *HTT*. The residual age at motor onset was calculated for each participant by subtracting the expected age at onset given pure CAG length from the observed age at motor onset (GeM-HD Consortium, 2015, 2019). Expected age at onset was estimated using pure CAG length based on the Langbehn equation that was derived for 41-55 CAGs (Langbehn et al., 2004), For 38, 39 and 40 CAGs we used 65, 62 and 59 years, respectively, extrapolating from Langbehn model. Observed age at motor onset was determined as follows: where onset was classified as motor, oculomotor or mixed, the rating clinician’s best estimate for motor onset was used (sxrater). For non-motor onsets, or where the clinician’s onset estimate was missing, the motor component of the clinical characteristics questionnaire was instead used (ccmtrage; (McAllister et al., 2021)). The Registry population was stratified by residual age at onset and 250 (∼ 4%) at each extreme of the distribution selected for analysis (exact samples depending on DNA availability) along with 7 technical exome-sequencing controls of different CAG repeat lengths and residuals of zero. Following targeted *HTT* CAG repeat sequencing using MiSeq (Figure 2), we determined accurate residual ages at motor onset for the Registry population using sequenced pure CAG lengths. These corrected residuals were used in all subsequent analyses.

The second cohort (N=238) was taken from CAG expansion carriers in the global (mainly US) Predict-HD study (total N=1069) that enrolled healthy at-risk *HTT* mutation carriers and prospectively followed then to clinical onset over ten years (Paulsen et al., 2008). Extremes were selected in two ways: first, phenotypic extremes for Total Motor Score (TMS) and Symbol Digit Modalities Test (SDMT) were selected from the PREDICT-HD database. The two variables were chosen because they represent the two important HD domains of motor (TMS) and cognitive (SDMT), and data was available for almost all individuals. Extremes were required to persist across the first and third visits for the participants in order to help ensure consistency. Locally weighted scatterplot smoothing (LOESS; (Cleveland et al., 1992)) was used to predict each variable using CAG, Age at study entry, and their interaction, with the predictors as a group being an index of HD progression (Zhang et al. 2011). LOESS residuals were computed, which reflect an individual’s deviation from the value that is expected based on their CAG/Age combination. For each visit, individuals were ranked by the residuals, and those who were consistently in the top or bottom 20% were selected as “consistent phenotypic extremes” for the exome analysis (N=117, TMS group; N=85, SDMT group).

Secondly, we selected a cohort with predicted early or predicted late onset HD. For those subjects who phenoconverted to Diagnostic Confidence Level 4 (DCL4) during the course of the study, the age at DCL4 was available as a proxy for age at onset. Using the PREDICT-HD database, extremes of expected onset were selected based on the relationship between the CAG-Age Product (CAP; (Zhang et al., 2011)) and 10-year survival from motor diagnosis (DCL = 4). CAP was computed as CAP = Age at study entry × (CAG − 33.66), and the predicted 10-year survival was the restricted mean survival time (RMST) computed for each participant. RMST was based on the machine learning method of random survival forest (RSF; (Ishwaran et al., 2008)) using progression in clinical variables to predict time to motor diagnosis. For each participant with two or more time points, the intercept and slope over time of Total Motor Score, Symbol Digit Modalities Test, Stroop interference, and total functional capacity, were estimated using ordinary least squares. Then the intercepts and slopes were used as predictors of time to DCL = 4 in the RSF. The RMST was computed by estimating the survival curve for each subject based on their predictor profile, and then taking the area under the curve up to 10 years. RMST is the “survival expectancy” (non-diagnosis expectancy) when we follow up a cohort from study entry to 10 years (Royston and Parmar, 2013). The participants were then subdivided into 20 groups of 53-54 by CAP score and the mean RMST determined for each group. For each participant, the absolute difference between the individual RMST and the RMST of their group was calculated.

Among the ∼6% extremes, 63 relatively earlier RMST and 56 relatively later RMST had DNA available and were included in whole-exome sequencing (N=119 total).

Pure CAG repeat lengths in expanded *HTT* alleles were initially determined by standard PCR-fragment length analysis in Predict-HD. To determine glutamine-encoding *HTT* repeat tract sequences, whole-exome sequencing data were extracted and manually assessed. Although it was not possible to determine full allelic structures as read lengths were 75 bp, phase could be determined for non-canonical alleles as unexpanded CAG lengths were short enough to be effectively captured (Table S3). Using this information, pure CAG repeat lengths were adjusted and used in all analyses. The presence or absence of extra non-canonical CAACAG or CAA or CAC triplets was used as a covariate in analyses

#### Ethical Approvals

Ethical approval for Registry was obtained in each participating country.

Investigation of deidentified Predict-HD subjects was approved by the Institutional Review Board of Partners HealthCare (now Mass General Brigham). Participants from both studies gave written informed consent. All experiments described herein were conducted in accordance with the declaration of Helsinki.

#### HD iPSC model

Human induced pluripotent stem cells (iPSCs) were generated previously from a human fibroblast line with an expanded *HTT* allele initially containing 109 pure CAG trinucleotides (Q109) (HD iPSC Consortium, 2012). Q109N1 and Q109N5 refer to clonal lines independently generated from fibroblasts from the same individual.

#### Generation of cell lines (see Figure S5)

*FAN1 knock-out*: Two guide RNAs (gRNAs) targeting exon 2 of FAN1; 5’-CTGATTGATAAGCTTCTACG**AGG**-3’ and 5’-GCACCATTTTACTGCAAACG**GGG**-3’ were designed on DESKGEN cloud (www.deskgen.com) to produce a 95 bp deletion. crRNA and tracrRNA-ATTO-550 (IDT) were combined in nuclease-free duplex buffer (IDT), annealed (95°C, 2 minutes), combined with Cas9 (IDT) and incubated (RT, 20 minutes) to form a ribonucleoprotein (RNP). iPSCs were nucleofected with both RNPs using the 4D-Nucleofector and P3 Primary Cell 4D-Nucleofector X Kit (Lonza). After 24 hours, iPSCs were sorted on the FACS ARIA Fusion to obtain the top 10% of cells, which were plated as single cells. After 7 days, individual colonies were manually dislodged and plated into single wells of a 96-well plate, which, after 7 days, were passaged into replicate plates using Gentle Cell Dissociation Reagent (STEMCELL Technologies). For screening DNA was extracted using QuickExtract (Cambio) (RT 10 minutes, 65°C 6 minutes, 95°C 2 minutes) and PCR amplified using two primer pairs amplifying exon 2 of FAN1; FAN-KO, 5’-CCTGTGTTTTATTGCTCAGAACA-3’ and 5’-CATTTCATCAAGGTGCCGGT-3’ and FAN1-T7, 5’-TCAGAGTTCGCTTTTCCCCT-3’ and 5’-GATGCTAGGCTTCCCAAACA-3’. Amplicons were visualised on a 1.5% agarose gel on the Geldoc XR system (Bio-Rad). Sanger sequencing was used to confirm successful editing.

*D960A variant;* The FAN1 variants D960A were introduced by utilising cellular homology directed repair (HDR). A single gRNA sequence (5’-AGGGGGCCTCCCCGACCTGG**TGG**-3’) and a 122 bp repair template containing the desired edit (5’-AGGaGGCCTCCCCG**c**CCTGGTcG-3’) provided a template for HDR. The HDR repair template contained two silent mutations (5’-AGG**a**GGCCTCCCCGcCCTGGT**c**G-3’), one of which was in the PAM sequence to prevent further excision, and introduced restriction sites enabling efficient screening. Nucleofection was carried out as above. For screening DNA was extracted using QuickExtract and PCR amplified using two primer pairs: FauI-D960A, 5’-TCACGAGGGAAGTGGCTAAC-3’ and 5’-GCCAACAGCCACTCAAGAAATG-3’ and StuI-D960A, 5’-TCACGAGGGAAGTGGCTAAC-3’ and 5’-CACAGAATACAGCAGGAGTGATG-3’. Restriction digest was performed with FauI or StuI in CutSmart Buffer (NEB). Amplicons were separated on a 1.5% agarose gel and visualised on the Geldoc XR system (Bio-Rad). Sanger sequencing was used to confirm successful editing.

## Method details

### Sequencing of *HTT* exon 1 repeat locus using MiSeq

We utilised a MiSeq-based *HTT* sizing protocol as described elsewhere (Ciosi et al., 2018, 2019; GeM-HD Consortium, 2019). Briefly, low-passage lymphoblastoid DNA was equilibrated using PicoGreen™ to 4 ngµL^−1^. Libraries were prepared in 384-plate format using allele specific primers targeting the *HTT* CAG and CCG repeats. Each PCR reaction contained 1.2 µL of each the forward and reverse primer (10 µM), 10 ng patient DNA, 0.55 µL nuclease-free water, 0.15 µL TaKaRa LA Taq® polymerase (RR02AG, TaKaRa), 7.5 µLTaKaRa GC Buffer II and 2.4 µL dNTPs and. PCR conditions used were 94^0^C for 5 min, followed by 28 cycles of 94^0^C (60 s), 60^0^C (60 s) and 72^0^C (120 s), followed by 10 min at 72^0^C. 2 µL nuclease-free water was used in place of human DNA for negative controls. Library clean-up consisted of two AMPure XP SPRI bead (Beckman Coulter, A63881) steps, performed using Beckman Coulter’s recommended parameters. The first clean-up used 0.6X bead concentration, and the second used 1.4X. Libraries were checked using a Bioanalyser (Agilent) with a high sensitivity DNA chip (Agilent, 5067-4626). Libraries were sequenced using a 600-cycle MiSeq v3 reagent kit (Illumina, MS-102-3003), running with 400bp forward and 200bp reverse sequencing. The sequencing parameters are described in (Ciosi et al., 2018).

### *HTT* genotyping and allele structure determination

MiSeq were genotyped and analysed as per (Ciosi et al., 2019) using the Scale-HD pipeline (v0.322; https://github.com/helloabunai/ScaleHD). Aligned sequence was also manually checked using tablet (Milne et al., 2013) in case rare alleles without known ref-seq were identified, which occurred in four instances on three alleles: (CAG)_n_**CAC**(CAG)_3_**CAA**CAG, (CAG)_n_(**CAA**)_2_CAG & (CAG)_n_(**CAA**)_3_CAG. Two Registry-HD samples failed MiSeq sequencing; for these samples, and for all Predict-HD samples, reads were extracted from whole-exome data using samtools (Li et al., 2009) using region 4:3073000-308000. Reads were manually assessed for presence of atypically interrupted *HTT* alleles. However, while it was possible to dephase most atypical interruptions by inference, it was impossible to determine the full structure and length of the *HTT* allele due working with short reads from exomes (75bp). Therefore, these individuals are not included in the analyses of the effects of non-canonical repeat structures on age at onset. A summary of Predict-HD interruption status is available in Table S3.

### Whole exome sequencing

For the Registry-HD cohort (N=507), sequencing was performed at Cardiff University. Whole-exome libraries were generated using TruSeq® rapid exome library kits (Illumina, 20020617) according to Illumina protocols (https://emea.support.illumina.com/downloads/truseq-rapid-exome-library-prep-reference-guide-1000000000751.html). DNA was obtained directly from BioRep, Italy, derived from low-passage lymphoblastoid cells. DNA was equilibrated using PicoGreen™ (P7589). Sequencing was performed on a HiSeq 4000. Libraries were pooled in equimolar amounts in groups of 96 and run over 8 lanes on a HiSeq 4000 patterned flowcell. Clustering used Illumina ExAmp regents from a HiSeq 3000/4000 PE cluster kit (Illumina, PE-410-1001) on a cBot system. Sequencing used a 2×75bp end run with a HiSeq 3000/4000 SBS kit for 150 cycles (Illumina, FC-410-1002).

For the Predict-HD participants, an in-solution DNA probe based hybrid selection method that uses similar principles as the Broad Institute-Agilent Technologies developed in-solution RNA probe based hybrid selection method (for details please see (Gnirke et al., 2009; Fisher et al., 2011)) was used to generate Illumina exome sequencing libraries from blood DNA. The Illumina exome specifically targets approximately 37.7Mb of mainly exonic territory made up of all targets from Agilent SureSelect All Exon V2, all coding regions of Gencode V11 genes, and all coding regions of RefSeq gene and KnownGene tracks from the UCSC genome browser (http://genome.ucsc.edu). Pooled libraries were normalized to 2nM and denatured using 0.2 N NaOH prior to sequencing. Flowcell cluster amplification and sequencing were performed according to the manufacturer’s protocols using the HiSeq 2500. Each run was a 76 bp paired-end with a dual eight-base index barcode read. Data were analyzed using the Broad Picard Pipeline which includes de-multiplexing and data aggregation.

### Sanger sequencing

All FAN1 variants identified by exome sequencing were confirmed by Sanger sequencing in the Registry-HD sample. Amplifications for sanger sequencing were performed using MyTaq™ (Bioline, BIO21127) as per the Bioline’s guidelines for PCR amplification. Briefly, a master mix containing 10 μL of 5x MyTaq reaction buffer, 0.5 μL of each forward and reverse primer (10 μM) and 26 μL of nuclease free water. 2 μL of each DNA template was added (∼10-30 ngμL-1). The programme was 95^0^C (2 min), followed by 30 cycles of 95^0^C (30s), 55^0^C (30s) and 72^0^C (60s), finishing with 72^0^C (5 min). PCR reactions were cleaned up using QIAquick PCR purification kits (28104, Qiagen) using the provided protocol for obtaining higher concentration purified samples. 1.5 μL of the relevant sequencing (Seq) primer (25 μM) was added to each purified sample (Table S9). Sequencing was performed externally using the Eurofins Genomics LIGHTRUN service.

### HD iPSC culture

iPSCs were cultured on Geltrex-coated plates (37 °C, 1 hour) (Life Technologies) in Essential 8 Flex medium (Life Technologies) under standard culturing conditions (37 °C, 5% CO_2_). iPSCs were passaged every 3-4 days using ReLeSR (STEMCELL Technologies) and seeded at a density of 1:12. Full media changes were performed every 1-2 days.

### HD iPSC differentiation

iPSC colonies were dissociated into a single cell suspension using Accutase (Life Technologies), seeded into 12 well plates coated with Growth Factor Reduced Matrigel (0.5 ng/mL) (BD Biosciences) and cultured in iPSC medium until the cells reached ∼80% confluency. iPSCs were differentiated to forebrain neurons using protocols adapted from (Telezhkin et al., 2016; Smith-Geater et al., 2020).

*NPC differentiation*: iPSCs were induced into NPCs using Advanced DMEM/F-12 (ADF) (Life Technologies) supplemented with 1% Glutamax (Thermo Fisher), 1% Penicillin/ Streptomycin (5000U/5000 μg) (Gibco), 2% MACS neurobrew without retinoic acid (Miltenyi), 10 µM SB431542 (Miltenyi), 1 µM LDN-193189 (StemGent) and 1.5 µM IWR-1-endo (Miltenyi) up until day 8, upon which SB was omitted from the medium and 25 ng/ml Activin A (PeproTech) was added. Full media changes were performed daily up until day 16. Day 16 NPCs were passaged into plates coated with Poly-D-Lysine (Thermo Fisher) and Growth Factor Reduced Matrigel. Cells were fed with SJA medium consisting of ADF with 1% Glutamax, 1% Penicillin/ Streptomycin, 2% MACS neurobrew with retinoic acid, 2 μM PDO332991 (Bio-Techne), 10 μM DAPT (Bio-Techne), 10 ng/ml BDNF (Miltenyi), 0.5 μM LM22A4 (Bio-Techne), 10 μM Forskolin (Bio-Techne), 3 μM CHIR 99021 (Bio-Techne), 0.3 mM GABA, 1.8 mM CaCl_2_ (Sigma-Aldrich) and 0.2 mM Ascorbic acid (Ascorbic Acid). After 7 days in SJA medium, cells were fed with SJB medium consisting of equal amounts of ADF and Neurobasal A (Life Technologies) with 1% Glutamax, 1% Penicillin/ Streptomycin, 2% MACS neurobrew with retinoic acid, 2 μM PDO332991, 10ng/ml BDNF, 3 μM CHIR 99021, 1.8mM CaCl_2_ and 0.2 mM Ascorbic acid. After 14 days in SJB medium, cells received half media changes every 3-4 day with BrainPhys Neuronal Medium (STEMCELL Technologies) supplemented with 1% Penicillin/ Streptomycin, 2% MACS neurobrew with Vitamin A and 10ng/ml BDNF.

### Lymphoblastoid cell line culture

Lymphoblastoid cell lines from individuals with HD were cultured under standard conditions (www.coriell.org) in RPMI-1640 Glutamax (Thermofisher) supplemented with 15% fetal bovine serum and 1% penicillin/streptomycin, and passaged three times per week.

### PCR-based fragment size analysis of *HTT* exon 1 repeat locus

DNA was extracted using the QIAamp DNA Mini Kit (QIAGEN), following manufacturer’s instructions. A fluorescently labelled forward primer FAM huHTT 1F: 5’-6-FAM-ATGAAGGCCTTCGAGTCCCTCAAGTCCTTC-3’ and a reverse primer: 5’-GGCGGCTGAGGAAGCTGAGGA-3’ were used to PCR amplify the CAG repeat region of *HTT*. Cycling conditions were as follows: initial denaturation at 94°C for 90s, followed by 35 cycles of 94°C for 30s, 65°C for 30s and 72°C for 90s and a final elongation at 72°C for 10mins. Resulting PCR products were sent for sizing with the GeneScan ™ 600 LIZ® dye Size Standard (Applied Biosystems) and run on the G3130xL Genetic Analyser (Applied Biosystems). Files were analysed using GeneMapper (Applied Biosystems), Fragman (Covarrubias-Pazaran et al., 2016) and AutoGenescan (https://github.com/BranduffMcli/AutoGenescan) and quantification was performed with a 10% peak height threshold applied (Lee et al., 2010).

### Western Blot

Cells were washed once in DPBS and lysed with RIPA buffer (Sigma-Aldrich) containing cOmplete™, EDTA-free Protease Inhibitor Cocktail Tablets (Merck). Protein samples were denatured at 70°C for 10 min in 4× NuPAGE LDS Sample Buffer (Life Technologies). 40 µg of cell protein extract per sample was separated on NuPAGE 4%-12% Bis-Tris gradient gels (Life Technologies) alongside PageRuler Plus Prestained Protein Ladder (Thermo Scientific) using MOPS SDS NuPAGE Running Buffer (Invitrogen) at 200 V for 50 minutes. Where purified proteins were run, 30 ng protein/lane were loaded and gels run in the same way. Gels were then transferred to methanol-activated Immobilon-P PVDF membranes (Sigma-Aldrich) using NuPAGE Transfer Buffer (Invitrogen) at 120 V for 45 minutes. The membrane was blocked in 5% milk in PBS-T and incubated overnight at 4°C with Anti-FAN1 (CHDI, sheep polyclonal, 1:1000) and Anti-β-Tubulin (UpState, mouse monoclonal, 1:10,000). Donkey anti-Mouse IgG Alexa Fluor 680 (Invitrogen, 1: 10,000) and IRDye^®^ 800CW Donkey anti-Goat IgG Secondary Antibody (LI-COR, 1:15,000) were used as secondary antibodies. Immunoblots were visualised with the Odyssey CLx Imaging System using β-Tubulin as a loading control.

### Cell Immunocytochemistry

iPSCs and neurons were fixed in 4% PFA for 15 minutes at room temperature. Cells were permeabilised in 0.1% Triton-X in PBS for 20 minutes at room temperature, and blocked in 3% BSA with 3% goat serum and 0.1% Triton-X. Primary antibodies used were OCT4 (Abcam, 1:100), CTIP2 (Abcam, 1:200) and MAP2 (Abcam, 1:500). Alexa Fluor goat anti-mouse 488 (Invitrogen, 1:400) and Alexa Fluor goat-anti rabbit 568 (Invitrogen, 1:800) were used as secondary antibodies.

### Mitomycin C survival assay

Lymphoblastoid cell lines (viability > 80%) were seeded in triplicate at 20000 cells/well in 12-well plates, treated once with mitomycin C (0 – 150 nM; SelleckChem), cultured for a further 7 days before viable cell counts determined by Trypan Blue staining (Countess II, Thermofisher). Each experiment was independently repeated three times.

### Protein purification and nuclease assay

Full-length Nus-His-FAN1 proteins were expressed in *E. coli* and partially purified in one step using cobalt-agarose (MacKay et al., 2010). Proteins were stored in 50 mM Tris-Cl pH 7.5, 270 mM sucrose,150 mM NaCl, 0.1 mM EGTA, 0.03% Brij-35, and 0.1 % β-mercaptoethanol.

dsDNA substrate containing a 5’ single-strand overhang with an IR-700 fluorescent label (5’ flap DNA) was generated by annealing three oligonucleotides (TOM112, TOM117, TOM122; based on (Rao et al., 2018)) at 95 °C for 10 min and cooling slowly to room temperature overnight. Nuclease assays were carried out by pre-incubating 10 nM Nus-His-FAN1 protein with 5 nM 5’ flap DNA in 9 μl reaction buffer (25 mM Tris-Cl pH 7.4, 25 mM NaCl, 0.1 mg/ml BSA, and 0.1 mM DTT) for 5 min on ice, then adding 1 μl MnCl_2_ (10 mM) and incubating at 37 °C to start reactions. Reactions were continued for 4 min before quenching with 10 μl stop buffer (1x TBE containing 12% Ficoll Type 400, 7 M urea, orange loading dye (LI-COR Biosciences) and 4 mM EDTA). Samples were denatured at 95°C for 5 min then run on a 15% TBE Urea gel (200 V, 1 h) before imaging on an Odyssey CLx (LI-COR Biosciences).

## Quantification and statistical analysis

### Alignment and variant calling

We utilised a standard Genome Analysis Toolkit (GATK, v3) best practices workflow for the alignment and variant calling of both sets of exomes (McKenna et al., 2010; DePristo et al., 2011; Van der Auwera et al., 2013). Reads were de-multiplexed, and adapters soft clipped using Picard (https://github.com/broadinstitute/picard). Alignment used BWA-MEM (Li and Durbin, 2009) to the hg19/GRCh37 genome assembly. Local insertion/deletion realignment was performed using GATK, and duplicate reads were marked and reads aggregated across lanes with Picard. Base quality scores were recalibrated using GATK’s base quality score recalibration (BQSR) technique. Germline SNPs were called with GATK’s haplotype caller (Poplin et al., 2017). Variant quality score recalibration (VQSR) was performed on both SNPs and insertion/deletion variants using GATK’s recommended parameters for exome sequencing.

### Exome quality control and annotation

Whole exome data was subject to a multi-step quality control pipeline (Figure S1A). Picard’s CollectHsMetrics function was used to assess target exome coverage; to be included in the study, exomes must have >=70% of the exome covered at 10x or greater. Per-sample mean genotype quality, mean depth and call rate were then determined using Hail (https://github.com/hail-is/hail). Exomes >3 standard deviations smaller than the mean of any of the three metrics were excluded for Registry-HD exomes. VerifyBamID (Jun et al., 2012) was used to detect contamination, and samples with a Freemix > 0.075 were excluded, as per the ExAC study (Lek et al., 2016). Where there were duplicate samples, the exome with the highest coverage was retained. Sex imputation used Peddy (Pedersen and Quinlan, 2017); samples with conflicting imputed and recorded sex were excluded. One individual with originally unknown sex was kept. Ancestry was estimated using Peddy by principal component analysis (PCA) against genomes from the 1000 genomes project phase 3 (1000 Genomes Project Consortium, 2015). Samples were excluded if they were either 1) predicted to have non-European ancestries by Peddy or 2) outside the primary cluster of European samples in Figure S2. First degree relatives were identified using Hail’s genetic relatedness matrix function, with a cut-off of 0.125. For each pair of related individuals, the individual with the most extreme uncorrected residual age at motor onset was retained.

Exomes underwent a multi-step annotation pipeline (Figure S1B). Exomes were annotated by gnomAD v2.1.1 (Karczewski et al., 2020), dbNSFP v4.0b2 (Liu et al., 2011, 2016) and Variant effect predictor (VEP) v95 (McLaren et al., 2016). Homozygotes were defined as having >=90% of reads as either the reference or the alternative allele, whereas heterozygotes were defined as having between 25%-75% of the reference or alternative allele. Loss-of-function (LoF) calls were defined as ‘HIGH’ impact calls by VEP, and non-synonymous damaging calls as either ‘HIGH’ or ‘MODERATE’ calls. Non-synonymous damaging calls included LoF calls or NS calls with ≥20 CADD score. Hail was used for principal component analysis (PCA), which were used in downstream analyses. Baseline variant rates (BVRs) were determined for each exome as the total number of variants classes at various minor allele frequencies (MAFs).

683 exomes passed quality control. After annotation, we identified 311,960 high-quality (≥75% call rate and ≥98.50 variant quality score recalibration) non-reference variants (Figure S3), with a mean of 35,808 variants per individual. There were 150,139 different non-synonymous variants (moderate and high impact variants) in our cohort with a mean of 12,700 such variants per individual, similar to previously reported population frequencies (Figure S3; (Lek et al., 2016)). Two groups of exomes were created for downstream analyses (Figure S1A and Table S2). First, a continuous phenotype group containing all those with known age at motor onset and calculable age at onset residual (N = 558; 95 Predict-HD, 463 Registry-HD). Second, a larger dichotomous group divided into extreme phenotypes (N = 637; 216 Predict-HD and 421 Registry-HD): individuals with early onset relative to CAG length (< −5 years age at motor onset residual), predicted early onset and/or worst 5% in TMS/SDMT (N = 315) and late onset relative to CAG length (> +5 years age at motor onset residual), predicted late onset and/or best 5% in TMS/SDMT (N = 322).

### Association analyses of rare variation

Coding variants that modify phenotype are likely to be deleterious to protein function. Therefore, our primary association analyses used variants meeting either of the following two criteria: (1) Loss-of-function variants (frameshifts, start/stop lost, premature stop codons, splice donor/acceptor variants). (2) Non-synonymous variants with a CADD-PHRED score ≥ 20 (i.e. the 1% predicted most damaging in the human genome) (Kircher et al., 2014). Additionally, variants were required to have minor allele frequency (MAF) < 0.01, as defined by the European cohort of gnomAD (v2.1.1) (Karczewski et al., 2020). To be included, variants required a call rate ≥ 75% and variant quality score recalibration (VQSR) ≥ 98.50. Secondary analyses were performed on loss-of-function and all non-synonymous variants, separately.

Given that coding variants of interest are individually rare, we collapsed qualifying coding variants on genes in the exome and tested each gene with at least 10 variants (N = 3912 genes (logistic group); N = 3198 genes (linear group) with at least 10 variant calls for association with residual HD onset using the optimised sequence kernel association test, SKAT-O (Wu et al., 2011; Lee et al., 2012b). SKAT-O combines elements of both a burden test and a Sequence Kernel Association Test (SKAT), and therefore does not assume that all variants in a gene have the same direction of effect. The sample with a dichotomous phenotype (N=637) were analysed *via* logistic regression, with late onset individuals, predicted late onset individuals and the best performing SDMT and/or TMS performers coded as 0 and early onset individuals, predicted early onset individuals and worst SDMT/TMS performers as 1. Those with a known quantitative age at onset residual (N = 558), were analysed *via* linear regression. We included population principal components 1-5, the baseline variant rate, mean sample depth and study group (Registry-HD or Predict-HD) as covariates in each regression analysis. Baseline variant rate was calculated for each individual and represented the total number of variants observed in the exome that passed QC at the particular minor allele frequency/damaging filter being used. Multiple testing correction was performed via a Bonferroni correction for the number of genes tested in each analysis. A burden test was additionally run using the same cut-offs and covariates as SKAT-O on the logistic patient group (N=637) using a Wald logistic burden regression test, implemented in Hail.

### Statistical modelling/analysis for iPSC data

The primary outcome measure was change in modal CAG from its initial value. Secondary analyses look at changes in expansion and instability index. All these outcome measures are zero when time is zero, requiring regression models without intercepts to be fitted. These are detailed below.

*D960-D42 iPSC data*: Data consisted of 3 wild-type, 3 FAN1^WT/D960A^ heterozygous, 3 FAN1^D960A/D960A^ clonal lines and 2 FAN1^-/-^ knock-out lines. Each clonal line was cultured in triplicate wells that remained independent from one another for the duration of the experiment and were repeatedly measured at different time points. Observations are therefore correlated if they are taken from the same line and/or well and it is important that statistical analyses take these correlations into account. This was done by performing mixed effects linear regression using the lmer() function in R, fitting random effects for the variation in rate of change of outcome between lines and wells. The models fitted were:

m0 <- lmer(change ∼0 + time + (0+time|line) + (0+time|well))
m1 <- lmer(change ∼0 + time + time:geno + (0+time|line) + (0+time|well))
The significance of different genotypes on rate of change of outcome is calculated by anova(m1,m0)

Genotype was initially coded as a four-level factor: 1=WT/WT, 2= 960A/WT, 3=960A/960A, 4= -/-, giving a 3df test. Post-hoc analyses on the pattern of genotype differences were performed by fitting models with restrictions on the genotype effects and comparing (via anova) to the general 3df model. Estimates of expansion rates for each genotype were produced by the R command m2<- lstrends(m1, ∼geno, var = “time”), and post-hoc pairwise comparisons of genotype effects using the Tukey method were obtained by the command pairs(m2).

*FAN1 knockout (neuronal data)*: Data consisted of 5 lines of N1-FAN1^+/+^ and 4 lines of N1-FAN1^-/-^. A separate, independent well was taken from each line at each time point. So, observations are correlated only through shared line, not through shared well. Effect of genotype (+/+ vs -/-) on outcome was tested by fitting the following two zero intercept mixed effect linear models:

m0 <- lmer(change ∼ 0 + time + (0 + time|line))
m1 <- lmer(change ∼ 0 + time + time:geno + (0 + time|line))
Again, the effect of genotype on the rate of change of the measures is tested by anova(m0,m1)

*FAN1 knockout (iPSC data)*: Data consisted of 7 WT (FAN1^+/+^) lines, and 6 FAN1 knock-out (FAN1^-/-^) lines in this experiment. Three of the FAN1^+/+^ and two of the FAN1^-/-^ lines were cultured in triplicate wells, the remainder in single wells. Wells remained independent from one another for the duration of the experiment and were repeatedly measured at different time points. Effect of genotype on the rate of change of the outcome phenotype was analysed using the same regression models that were used for the D960A-D42 iPSC experiment.

*D960A neural precursor data*: Data consist of 1 line each of WT/WT, D960A/WT and D960A/D960A. A separate, independent well was taken from each line at each time point. So, observations are correlated only through shared line, not through shared well. Effect of genotype on rate of change of outcome was tested by fitting the same zero intercept mixed effect linear models as were used to analyse the FAN1 knockout neuronal data.

### FAN1 Structure

FAN1 structure annotation used FAN1 in complex with DNA (Wang et al., 2014). This structure (PDB ID: 4RI8) contained residues 370-1017, the UBZ and subsequent unstructured domain having been removed, additionally two small loop sections were missing and added to the model using the homology model tools within MOE (Molecular Operating Environment).

